# Perceptual similarity judgments reflect one’s own discrimination capacities

**DOI:** 10.1101/2024.06.13.598769

**Authors:** Ali Moharramipour, William Zhou, Dobromir Rahnev, Hakwan Lau

## Abstract

Perceptual similarity is a cornerstone for human learning and generalization. However, in assessing perceptual similarity between two stimuli, it is not well-defined which features or combinations of features one should focus on. The problem has accordingly been considered ill-posed, implying a lack of a clear ground-truth basis for similarity judgments. We hypothesized that the stimuli rated as subjectively similar are those that are, in fact, more challenging for oneself to discern in practice, in near-threshold settings (e.g., psychophysics experiments). Therefore, an individual’s perceptual discrimination capacities can serve as a quasi-objective ground truth for determining whether two stimuli should be judged as similar. To test this idea, we asked participants to rank the similarity of face pairs and also separately measured their discrimination capacity for those pairs. We found a positive association between perceptual discrimination capacity and subjective perceptual dissimilarity, with this association being specific to each individual. The results indicate that perceptual similarity judgments reflect and predict one’s own perceptual discrimination capacities. Overall, although subjective, similarity judgments are normative in that they are grounded in one’s perceptual capacities, which can be measured through a psychophysical task with objective responses.

## 1. Introduction

Subjective perceptual similarity has long been a topic of research in behavioral science. Previous studies have explored various factors modulating similarity judgments, such as the effects of knowledge and expertise, contextual cues, and the order of presenting the stimuli (Medin et al., 1993; Shepard, 1964; Smith, 1989; Smith & Heise, 1992; Tversky, 1977). Different theories and quantitative models of similarity have also been proposed (Nosofsky, 1984; Shepard, 1987; Smith, 1989; Tversky, 1977). For example, Roger Shepard famously formulated the universal law of generalization, stating that perceived similarity decreases exponentially as stimuli become more distant from each other in the mind’s internal representation of them (Shepard, 1987). Intriguingly, recent research has demonstrated that the exponential similarity decay, coupled with a signal detection theory, can also effectively capture observations in visual working memory (Schurgin et al., 2020). There is also a rich history of studies utilizing similarity judgments, in combination with multidimensional scaling, to uncover the underlying perceptual dimensions of stimuli (Borg & Groenen, 2005; Hebart et al., 2020).

Similarity judgments are subjective, in that it is up to the subject to report how they feel about the stimuli. Accordingly, some researchers have argued that similarity judgments may reflect key aspects of conscious perception (Clark, 2000; Lau et al., 2022; Malach, 2021; Moharramipour & Lau, 2024; Rosenthal, 2010; Tallon-Baudry, 2022; Zeleznikow-Johnston et al., 2023). However, the essentially subjective nature of these judgments also led to the well-known critique that similarity is perhaps an ill-posed problem: there is, in a sense, no objective ground-truth as to how similar two things really are (Goodman, 1972; Medin et al., 1993). For example, Joe Biden may look more similar to Hillary Clinton than to Barack Obama, with respect to skin color. However, if we focus on gender-related facial features, Joe Biden may look more similar to Barack Obama. From the outset, it is unclear which visual features one should focus on. This presents a challenging obstacle to understanding the processes underlying similarity judgments, as mechanistic explanations of perception often rely on characterizing the observer as performing optimal inference, given existing constraints (Rao, 1999; Shen & Ma, 2016).

Following previous theoretical work (Lau et al., 2022), we hypothesize that subjective similarity judgments may be normative and rational, in the sense that they are made systematically based on our own perceptual abilities: stimulus pairs judged to be more similar are, in fact, more challenging for oneself to discern in practice. If one judges two stimuli to be highly dissimilar, and yet fails to distinguish them in psychophysical tasks, the said similarity judgment can be regarded as ‘incorrect’ in a meaningful sense.

Revisiting the above example of how subjectively similar faces are, the idea is that such a similarity judgment would rely on a combination of relevant features in a way that maximally differentiates the faces. Specifically, for this combination to be optimal, features should be weighted according to their perceptibility to the observer. In other words, this process is not only about physical stimulus itself, but rather, it reflects one’s own perceptual abilities.

The above is a non-trivial prediction, because an alternative hypothesis is that subjective similarity ratings may be made based on whatever visual features happen to be more salient, depending on one’s fluctuating attentional states, or arbitrary preferences that are not necessarily related to one’s own performance in near-threshold psychophysical tasks. This alternative hypothesis is not implausible given that ‘error’ feedback is generally never given to subjects, to ‘correct’ them or train them, as they make these similarity ratings somewhat freely.

To test our hypothesis, we quantified the degree of subjective perceptual similarity between stimulus pairs by having participants freely rank similarity, without being given specific criteria, across a stimulus set (Figure 1B; subjective similarity judgment task). We also assessed participants’ perceptual discrimination capacity between the pairs. The stimulus pairs may be so obviously dissimilar that discriminating between them is just too easy (i.e., performance under normal conditions would be at the ceiling). To address this problem, we proposed to use a psychophysical method to measure such discrimination capacity near the perceptual threshold. We measured participants’ discrimination performance within the morph set that spans between the two stimuli (Figure 1C; near-threshold discrimination task). With this, we quantified the number of just-noticeable-differences (#JNDs; see legends of Figure 1C for explanation) between a pair. The #JNDs reflects perceptual discrimination capacity, with higher values indicating greater capacity. We used faces as stimuli in our study due to their highly complex (i.e., high-dimensional) nature, and the fact that these are naturalistic stimuli commonly encountered in everyday life. In subsequent sections, we use the notion “dissimilarity” instead of “similarity”, so the hypothesized association with discrimination capacity is positive.

**Figure 1.**
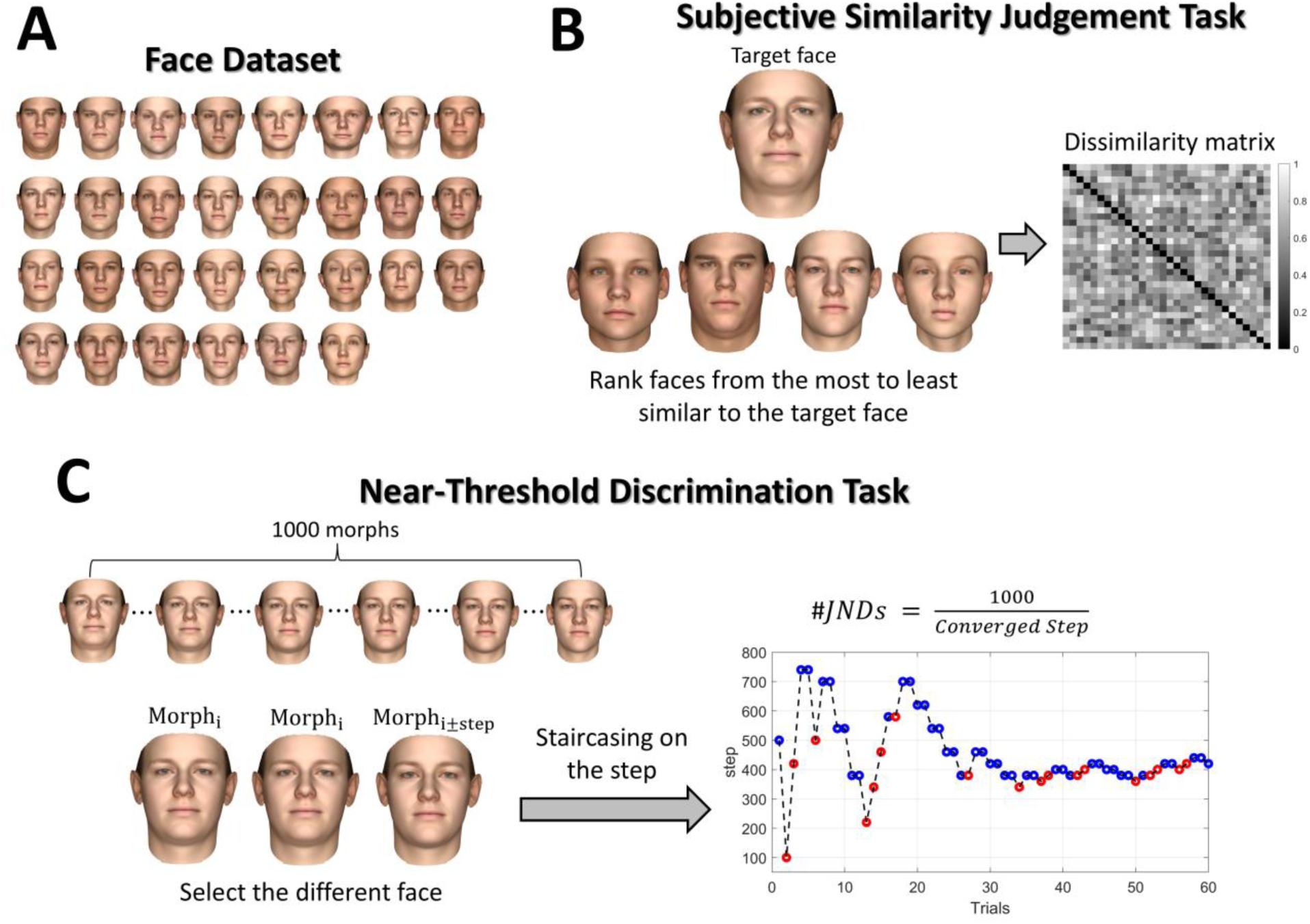
Experimental tasks for estimating subjective perceptual dissimilarity and perceptual discrimination capacity. **(A)** Illustration of 30 faces used in the present study. **(B)** The subjective similarity judgment task for estimating the level of subjective perceptual dissimilarity between face pairs for each participant. A target face on top and four other faces (candidates) on the bottom were shown in each trial. Participants were instructed to rank the candidate faces from the most to least similar to the target face by clicking on them in order. Then, a 30×30 dissimilarity matrix was computed from the participant’s responses across trials, with the value in each cell of the matrix indicating the level of subjective dissimilarity between a face pair. **(C)** The near-threshold perceptual discrimination task for measuring each participant’s discrimination capacity between two faces. One thousand morphs were created as intermediate transitions between two faces (based on a computational face model; see Methods for details). In each trial, three faces were shown simultaneously to the participants. Two were identical, and one was different from the other two by a certain degree (number of morph steps within the 1000-morph series). Participants were instructed to click on the face that was different from the other two. Task difficulty was maintained by titrating the number of morph steps needed for the different face to be barely detectable, using a standard 1-up-2-down staircase method. The converged (i.e., stabilized) staircase value indicates the number of morph steps required to maintain near-threshold performance (71% correct); thus, it reflects the just-noticeable-difference (JND). Because these morph steps come from a series of 1000 morphs between a face pair (e.g., any two faces in 1A), if e.g. JND = 250 morph steps, we can also describe the two faces concerned as being 4 JNDs apart from each other. This general notion of the number of JNDs (#JNDs) between face pairs, which is just the total number of morph steps (1000) divided by the measured JND, allows us to describe the psychophysical discriminability between two faces, free from the non-standard physical unit of ‘morph steps’ (which depends on the arbitrary specifics of the morphing procedure such as total number of steps used). Essentially, #JNDs indicates the perceptual distance between a face pair, in other words, how many JNDs are in between the face pair; thus, its higher value corresponds to a higher discrimination capacity.

In summary, we hypothesize that there is a correlation between perceptual discrimination capacity (in a near-threshold task with objectively defined correct responses) and subjective perceptual dissimilarity (as reflected by self-ratings of supra-threshold stimulus pairs). Specifically, perceptual discrimination capacities are higher in face pairs that are subjectively judged to be more dissimilar (Hypothesis 1). Further–and critically–we hypothesize that this association is specific to each individual, such that one’s subjective perceptual dissimilarity is better explained by one’s own perceptual discrimination capacity than others’ discrimination capacity (Hypothesis 2). This would support the idea that one’s perceptual capacity can serve as a quasi-objective ground truth basis for their subjective similarity judgment. An overview of the hypotheses, their corresponding tests, and potential interpretations of different outcomes is provided in Supplementary Table 1.

## 2. Methods

### 2.1. Ethics information

The study received approval from the Ethics Review Committee at RIKEN, complying with all their ethical guidelines. Informed consent was obtained from participants before the experiment, and in appreciation of their participation, they were compensated with 3500 yen (approximately 23 US dollars) per day (roughly 90 minutes).

### 2.2. Design

Twelve participants were recruited for the study. They initially performed the subjective similarity judgment task, twice over the course of two days. After all participants completed this task twice, a set of 24 face pairs was selected for examination in the near-threshold discrimination task. The criteria for selecting the face pairs are described in section 2.2.3. Then, all participants were invited back to perform the near-threshold discrimination task over two days on these 24 face pairs, with the pairs randomly split across the days.

Note that there were 48 sessions in total, across 12 participants. Each participant completed four sessions on different days, with each session taking more than 60 minutes. This provided us with sufficient data to perform our statistical analysis even at the individual level. The subjective similarity judgment task comprised 330 carefully designed trials to estimate the degree of subjective perceptual dissimilarity between all face pairs. Participants performed this task twice, and the resulting dissimilarity values were averaged to enhance robustness. The near-threshold discrimination task comprised 1440 trials (60 per face pair) to estimate the perceptual discrimination capacity between a systematically selected set of 24 face pairs. We also pre-registered that we would recruit more participants if the initial 12 participants did not satisfy our data-collection stopping rule described in section 2.3. However, since they met the stopping criterion, no additional participants were recruited. As discussed in section 2.3, this criterion ensured that the obtained statistics had sufficient precision to draw robust conclusions.

Note that this study was pre-registered and <u>peer-reviewed</u> at *Peer Community In (PCI)* as a Registered Report (Stage 1), with the <u>preprint</u> available at (Moharramipour et al., 2024). The study was conducted in accordance with the recommended Stage 1 protocol.

#### 2.2.1. Face data set

The basal face model (BFM) (Paysan et al., 2009) was used to select our face dataset and generate morphs between the faces for the near-threshold discrimination task. BFM is a widely used morphable model for generating graphical faces with two embedded vectors describing the shape and texture of the faces independently. We arbitrarily selected 30 faces from the BFM space, while ensuring a diverse set that includes faces positioned both close to and far from one another within the BFM space. The top three shape dimensions were assigned systematically from a cylindrical coordinate with a 2.5 SD radius, and the subsequent top 47 shape dimensions and top 50 texture dimensions were assigned randomly from a uniform distribution ranging between -1.5 and 1.5 SD. The remaining less important shape and texture dimensions were set to zero. Figure 1A shows the selected 30 faces for the study.

#### 2.2.2. Subjective similarity judgment task

In each trial, participants were presented with a visual arrangement consisting of one face positioned at the top (target face) and four other faces positioned at the bottom (candidate faces) (Figure 1B). Participants were instructed to rank the four bottom candidate faces based on their perceived similarity to the top target face by mouse-clicking on the faces in the order of most to least similar. Each clicked face immediately disappeared from the screen, and the trial ended after all candidate faces were clicked one by one. If participants failed to complete the trial within 30 seconds, the trial was skipped, and any ranking assigned was disregarded. The aim of this task was to estimate the level of subjective dissimilarity between each face pair and to construct a dissimilarity matrix (Figure 1B) for each participant by analyzing their ranking data across trials.

The level of subjective dissimilarity (dissimilarity value) between two faces was estimated by calculating the probability of one face being ranked lower than the rest of the faces when the other face was the target, as outlined in the following. The rankings given in all trials were segmented into sets of three, consisting of the target face and the combination of two of the four candidate faces (i.e., six sets per trial). Within each set, the face that ranked lower was marked as the odd face. Subsequently, the dissimilarity value between a face pair was determined by calculating the ratio of instances where one of the faces was marked as the odd face across all sets that included the face pair with either of them as the target face. Note that when calculating this ratio, we accounted for a non-tested set with an obvious outcome, where one of the faces repeats, by adding 0.5 to both the numerator and the denominator. This fundamentally prevented getting a dissimilarity value of zero, as only the diagonal value of the dissimilarity matrix should be zero.

The tuple of five faces displayed in each trial was strategically selected using the InfoTuple method (Canal et al., 2020). This method guaranteed that each trial offered informative data, thereby enhancing the estimation of the dissimilarity matrix. This essentially enabled achieving a robust estimation of the dissimilarity matrix over a smaller number of trials. The trial selection procedure was similar to the one used by Canal et al. (Canal et al., 2020) and comprised the following steps:

1. The tuple set for the first 30 trials was selected at random while ensuring that each face was selected once as the target face.
2. The dissimilarity matrix was calculated as described above, and a 5-dimensional metric multidimensional scaling (MDS) (Borg & Groenen, 2005) was applied to the dissimilarity matrix to find its embeddings.
3. A cycle of 30 trials, showcasing each face once as the target face, was selected by the InfoTuple method using the embeddings. The InfoTuple method selects the tuple that maximizes a mutual information estimate, which involves two entropy terms: intuitively, one term favors tuples whose rankings are uncertain given the current embeddings, while the other discourages inherently ambiguous tuples that are expected to remain uncertain even if the embeddings are revealed. So, it aims to select an informative tuple whose rankings are unknown but yet can be answered reliably (consistently). Please refer to the original paper for a detailed explanation and the mathematics of the InfoTuple (Canal et al., 2020).
4. The dissimilarity matrix was calculated from all data collected thus far, and the embeddings were updated by applying a 5D metric MDS to the dissimilarity matrix, using the previous embeddings as the initial seed.
5. Steps 3 and 4 were repeated for 10 iterations. We stopped after 10 iterations as the dissimilarity matrix and embeddings were expected to reach a relatively stable state at this point. As completing 10 iterations plus the initial burn-in iteration is lengthy and can be exhausting, participants were given a short break after every three iterations.

Lastly, the embeddings from the 5D metric MDS were used to recalculate the dissimilarity matrix by computing the Euclidean distances between the embedded face points. This process of reducing dimensionality to 5D denoises the original dissimilarity matrix, refining inaccuracies in some cells caused by insufficient data or noise in responses (i.e., response inconsistencies). This reconstructed dissimilarity matrix was then utilized in all the subsequent stages instead of the original dissimilarity matrix. Please see Supplementary Figure 1 for a schematic overview of the task design described above.

We also pre-registered that we would test using a 2D metric MDS to reconstruct the dissimilarity matrix. Even though there might be some information loss in using such a low-dimensional MDS, it may further denoise the matrix. We thought that focusing on only two primary components of the dissimilarity relations may potentially make their association with perceptual discrimination capacity more salient. This prediction was based on our pilot data (see supplementary materials), in which both Hypotheses 1 and 2 achieved higher significance (p-value < 0.05 for all participants) when a 2D rather than a 5D MDS was used.

We note that, in our algorithm, prior to applying the metric MDS, a nonmetric MDS was used to fill in missing values in the dissimilarity matrix (i.e., pairs with no ranking data). The missing cells were filled in by the Euclidean distances computed from the embeddings derived by the nonmetric MDS. This procedure was important in the initial iterations, in which there was a considerable number of missing cells. The metric MDS was then applied to this filled-in dissimilarity matrix. We did not use the nonmetric MDS directly because, unlike the metric MDS, it does not preserve the magnitude of dissimilarity between pairs.

Each participant performed the above task twice, each time on a different day (i.e., the full task, including 1+10 iterations, as described above, on each day). The average of the dissimilarity matrices obtained from each day formed the final dissimilarity matrix. To evaluate the reliability of these estimates, we assessed the within-participant correlation between the dissimilarity matrices derived from each day. As a reference, we compared it with the distribution of between-participant correlations computed by randomly correlating a dissimilarity matrix from a session in one participant with that of another participant. We expected that the within-participant correlation would be higher than the between-participant correlation.

In addition to the above approach for estimating the dissimilarity matrix, we pre-registered a machine-learning approach for exploratory analysis (i.e., that we would not use this method for our main hypothesis testing or the stopping criterion. See sections 2.3 and 2.4). This approach starts with random embeddings and iteratively updates them to minimize a loss function that penalizes incorrect similarity rankings derived from the embeddings. The loss is constructed using a sigmoid activation function as follows:

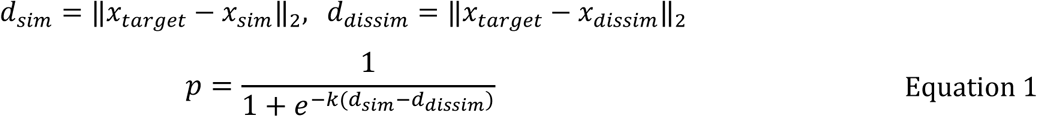

Where *x*_*m*_ represents the vector embedding of face *m*; *d*_*sim*_ and *d*_*dissim*_ are the Euclidean distances between a target face and a face ranked as more similar and a face ranked as less similar to the target face by the participant, respectively; *k* corresponds to the ranking difference (e.g., 2 for a face ranked first and a face ranked third) that sharpens the activation function *p* to put more emphasis on clearer similarity comparisons. Essentially, when *d*_*sim*_ is larger than *d*_*dissim*_, *p* takes a non-zero value, approaching one, indicating an error. Subsequently, *p* was used with a binary cross-entropy function to calculate the overall loss across all similarity comparisons as follows:

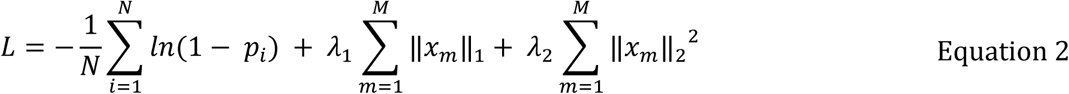

where *N* indicates the number of all segmented trio comparisons (with six trio segments in a trial); *λ*_1_ and *λ*_2_are the hyperparameters of L1 and L2 regularizations, which help to control the sparsity (i.e., suppressing unnecessary dimensions) and the scale of the embeddings, respectively. We used the Keras library in Python, with the Adam optimizer, to minimize the loss function, so that the resulting embeddings produce Euclidean distances between pairs that best reflect the participant’s similarity rankings.

Note that we did not use this machine-learning approach during the task (i.e., in our online application) because it was slower and required cross-validation to select appropriate hyperparameters. This approach was more sophisticated than our main approach, which simply involved estimating ranking probabilities and applying MDS to reduce dimensionality and thereby reduce noise. We pre-registered the machine-learning approach because we anticipated it could yield a more accurate estimate of the dissimilarity matrix. In our pilot study, using this machine-learning approach, we got similar results to those shown in Supplementary Figures 3 and 4 obtained with the main pre-registered approach.

The *λ*_1_and *λ*_2_values that effectively reduce embeddings dimensionally and scale, while maintaining accuracy on a validation set, are considered optimal as they result in more generalizable, thus more robust embeddings. The *λ*_1_and *λ*_2_were fixed at 0.00025, as they fell within the optimal range for all session data, identified through grid search sampling of *λ*_1_ and *λ*_2_ values. The embeddings were initialized with 10 dimensions. However, the optimal *λ*_1_ and *λ*_2_resulted in embeddings with 3 to 5 effective dimensions across all sessions, indicating that no more than five dimensions were required to represent how participants perceived the similarity among the 30 faces used in the study.

#### 2.2.3. Selection of the pairs for the near-threshold discrimination task

Following the completion of the subjective similarity judgment task twice by all 12 participants, 24 face pairs were systematically selected to be examined in the near-threshold discrimination task. Practical constraints (time limitations; it takes 4-5 minutes to complete the near-threshold discrimination task for a face pair) limited us to examining only a small subset of the pairs (24 out of 435 possible pairs). We expected a sample size of 24 pairs to be adequate for exploring our hypotheses. This was also supported by our pilot study (Supplementary Figure 3), which yielded reasonably robust results despite only 13 pairs being examined. Participants were reinvited to two more sessions to perform the near-threshold discrimination task on these 24 pairs, completing 12 pairs in each session.

While we examined only a limited subset of pairs, we carefully selected them to provide the most informative data for testing our hypotheses. Pairs with controversial subjective dissimilarity across participants (i.e., those in which participants showed substantial disagreement about their perceptual dissimilarity) were particularly promising candidates. If our hypotheses hold, these pairs should also exhibit controversial discrimination capacities across participants. Essentially, given the inherent noise in our methods for measuring dissimilarity values and discrimination capacities, any relation should be more readily detectable in pairs with larger deviations across participants. Accordingly, we selected 18 pairs with dissimilarity values that were controversial across participants. For comparison, we also selected 6 less-controversial pairs. Although these less-controversial pairs are not as informative, we included them to assess whether our measures have sufficient precision to detect effects (i.e., especially the individual-specific relationships; if Hypothesis 2 is true) when deviations across participants are smaller.

First, the dissimilarity matrix was z-normalized within each participant to ensure that its scale was consistent across participants. Subsequently, the median and the median absolute deviation (MAD) of the dissimilarity matrix were computed across participants, and the quantiles of the median values were derived. Within the first and the last quantiles, 3 pairs with the highest and 1 with the lowest MAD were selected. Additionally, 6 pairs with the highest and 2 pairs with the lowest MAD were chosen within the second and the third quantiles. This systematic selection ensured choosing 18 controversial and 6 less-controversial pairs that covered a diverse range in the group-averaged dissimilarity matrix.

#### 2.2.4. Near-threshold discrimination task

A series of 1000 equally spaced morphs was generated along the line connecting a face pair in the BFM space. In each trial, three faces were shown to the participants: two identical faces, randomly selected from the morph set, and a third different face spaced by a certain number of morphs (step value) from the identical faces (e.g., morph 200, morph 200, and morph 300: here the step value is 100). The faces were displayed simultaneously at the center of the screen next to each other, and their arrangement was randomized in each trial (Figure 1C). Participants were instructed to identify and click on the different face.

A staircase (Cornsweet, 1962) with a 1-up and 2-down protocol was applied to the step value (i.e., the number of morphs between the different and identical faces), initiating from a step value of 500. After each incorrect response trial, the step value was increased, and it was decreased after two consecutive correct trials. The magnitude of the change in the step value gradually decreased over trials, reaching a minimum change of 20 steps. The task was terminated after 60 trials, allowing precise convergence of the step value. The converged step value indicates the JND, the minimum degree of differences between the faces required to achieve near-threshold discrimination performance (71% correct response). The average of the steps achieved within the last 5 changes was defined as the converged step. Essentially, a small JND, for example, 125, indicates that the two questioned faces were quite distinct, involving 8 JNDs (i.e., 1000 divided by 125) between them. We used the notion of the number of JNDs (#JNDs) to quantify perceptual discrimination capacity. The #JNDs indicates the perceptual distance between a face pair, in other words, how many JNDs are in between the face pair for a participant. Therefore, its higher value corresponds to a higher discrimination capacity. The #JNDs was simply calculated as 1000 (i.e., total number of morphs) divided by the JND.

In a session, there were 12 face pairs to undergo the staircase procedure. The staircases for each of these face pairs were interleaved, progressing concurrently. There was a cycle of 12 trials, featuring each staircase once in a random order. The trials were time-constrained, requiring participants to respond within 8 seconds. If participants failed to respond within this time window, the trial was skipped and reintroduced at the end of the cycle. To encourage participants to perform to the best of their abilities, they were provided with feedback on their responses. A green circle was displayed on the different face (i.e., correct answer) and red crosses on the identical faces (i.e., wrong answers) after they provided their response. Since the session was lengthy, with a total of 720 trials, participants were given a short break after every 180 trials.

The trajectory convergence of a staircase could indicate the reliability of the estimated #JNDs. A staircase with a higher ratio of reversals in its later trials could be considered more reliable. Therefore, we assessed the ratio of reversals in the last 20 trials of each pair’s staircase and its statistics across participants. In an absolute ideal case, given our 1-up and 2-down staircase protocol, the ratio of reversals in the last 20 trials would be 0.6 at most.

### 2.3. Sampling

A set of criteria was pre-registered to exclude participants from the analysis. However, none of the participants met any of the exclusion criteria. The criteria were as follows: Those who did not complete all four experimental sessions and those who showed a lack of attentiveness to the task in any of the sessions. The lack of attentiveness in the near-threshold discrimination task was identified by non-converging staircases, indicated by a non-fluctuating increment in the step value over trials. Specifically, a session in which there were more than 4 (out of 12) staircases with less than three downs in their last 20 trials was considered bad, lacking sufficient attentiveness. None of the sessions from any participants met this criterion. In the subjective similarity judgment task, the lack of attentiveness was judged by comparing the consistency of responses between the first and second halves of the session. Specifically, if the correlation between the dissimilarity matrices estimated from each half fell below 0.2, the session was considered bad with inadequate attentiveness. This correlation ranged between 0.50 and 0.78 (mean ± SD= 0.65 ± 0.08) in all sessions, indicating good data quality.

A data-collection stopping rule was pre-registered to ensure that the collected data would be sufficient to draw robust conclusions. We pre-registered to recruit additional participants until the criterion was met, with a maximum sample size of 24 participants (see Supplementary Figure 2). However, because the criterion was satisfied with our initial 12 participants, no further participants were recruited, and the data collection was concluded. The criterion was as follows: data collection would be stopped if, in both Hypotheses 1 and 2, the width of the 95% confidence interval of the group-mean z-value was less than 1 (see section 2.4; one z-value statistic was computed at the individual participant level for each hypothesis). We also pre-registered that we would consider a hypothesis confirmed, if the group-mean z-value was significantly above zero, specifically, if the 95% confidence interval was above zero. Note that we set our stopping criterion independent of the significance testing and solely based on the precision of the effect (i.e., the confidence interval width). So that we are confident the effect is not being confirmed or rejected because of some extreme observations (Cumming, 2008; Lakens, 2014). Given our sample size scale, we expected a considerable effect to have a group-mean z-value of at least above 0.5. So, a minimally significant scenario would involve a group-mean z-value of 0.5 with a 95% confidence interval width less than 1. Considering this, we set our stopping criterion as the width of the 95% confidence interval being smaller than 1 to safely reject or accept the hypotheses.

### 2.4. Analysis

#### 2.4.1. Hypothesis 1

Spearman correlation coefficient and its p-value were computed between the dissimilarity values and #JNDs of the examined 24 pairs in each individual. The Spearman correlations were converted to z-values using the Fisher z-transformation (Fieller et al., 1957) to conduct group-level statistical tests. The distribution of the group-mean z-value was computed by bootstrapping, iterated 100,000 times, and then its 95% confidence interval was derived by obtaining the 2.5th and 97.5th percentiles of the distribution. The hypothesis was considered confirmed if this confidence interval was above zero.

The following statistics were also computed as complementary information: (1) the p-value and the Bayes factor of a t-test applied to z-values (BayesFactor Matlab package is used: https://zenodo.org/badge/latestdoi/162604707). Following standard convention, a Bayes factor value of 1–3 was taken as indicating weak evidence, 3–10 as moderate evidence, 10–30 as strong evidence, 30–100 as very strong evidence, and above 100 as decisive/extreme evidence for the hypothesis. Values below 1 were taken symmetrically as evidence against the hypothesis. (2) the p-value of a Fisher’s combined probability test, combining individual-level p-values (Brown, 1975). (3) a Bayesian posterior distribution of population prevalence (Ince et al., 2021) and its 95% highest posterior density interval, considering the p-value of 0.05 as the individual-level significance threshold. The Bayesian posterior distribution quantifies the likelihood of observing an effect in the population, given the number of participants tested in the study and the proportion who showed the effect at the individual level. (4) an equivalence test (Lakens et al., 2018), considering a z-value range of -0.5 to 0.5 as the smallest effect size of interest, to assess the significance of the null result (i.e., how confidently the effect can be considered non-significant). However, because no null results were observed, this test was not applied.

#### 2.4.2. Hypothesis 2

Each participant’s #JNDs were z-normalized to ensure that the #JNDs range was consistent across participants. This normalization was crucial, given that some participants exhibited generally higher #JNDs than others. Subsequently, a nonparametric permutation test was applied to each individual to assess the specificity of the relationship between their #JNDs and dissimilarity values, as follows:

1. A permutation set of #JNDs was constructed by randomly permuting the #JNDs across participants (i.e., for each pair, selecting the value in one of the participants at random), excluding the participant in question. Essentially, the permutation set simulated a new participant by mixing the existing participants.
2. The Spearman correlation coefficient was calculated between the permuted #JNDs and the dissimilarity values of the participant in question. It is noteworthy that with 12 participants and 24 pairs, there were an enormous number of possible permutations (i.e., 11^24 unique permutations), which ensured the construction of a reliable null distribution.
3. Steps 1 and 2 were repeated 100,000 times to derive the distribution of the Spearman coefficients. This distribution represents the null hypothesis distribution in which there is no individual specificity.
4. The actual Spearman coefficient between the #JNDs and dissimilarity values of the participant in question was tested against the null distribution, and the p-value, indicating the significance level, was derived. The z-value was also calculated by subtracting the actual Spearman correlation from the null distribution’s mean and then dividing it by the null distribution’s SD.

Then, similar to Hypothesis 1, the 95% confidence interval of the group-mean z-value was derived through bootstrapping, and if it was above zero, the hypothesis was considered confirmed. The complementary statistics, outlined in Hypothesis 1, were also computed for Hypothesis 2.

## 3. Results

### 3.1. Quality of measures

#### 3.1.1. Subjective perceptual dissimilarity

Participants performed the subjective similarity judgment task twice on two different days. Similarity judgments are expected to be relatively consistent within individuals. If, however, similarity judgments are not made consistently, changing from one session to the next, then this would automatically reject our hypothesis that similarity judgments are made normatively based on one’s perceptual capacities. Therefore, it was important to examine the consistency between participants’ first and second sessions of the subjective similarity judgment task. To this end, within-participant correlations—the Spearman correlation between the dissimilarity matrices derived from the first and second sessions of the same participants—were computed. These were then evaluated against the distribution of between-participant correlation obtained by randomly correlating a dissimilarity matrix from a session in one participant with that of another participant. As shown in Figure 2, the within-participant correlations were significantly higher than the between-participant correlations. The z-value of the within-participant correlation relative to the distribution of the between-participant correlation was highly above zero in all participants, with most (ten out of twelve participants) surpassing 2.3. This indicates that similarity judgments were consistent within each individual and also unique to them.

**Figure 2.**
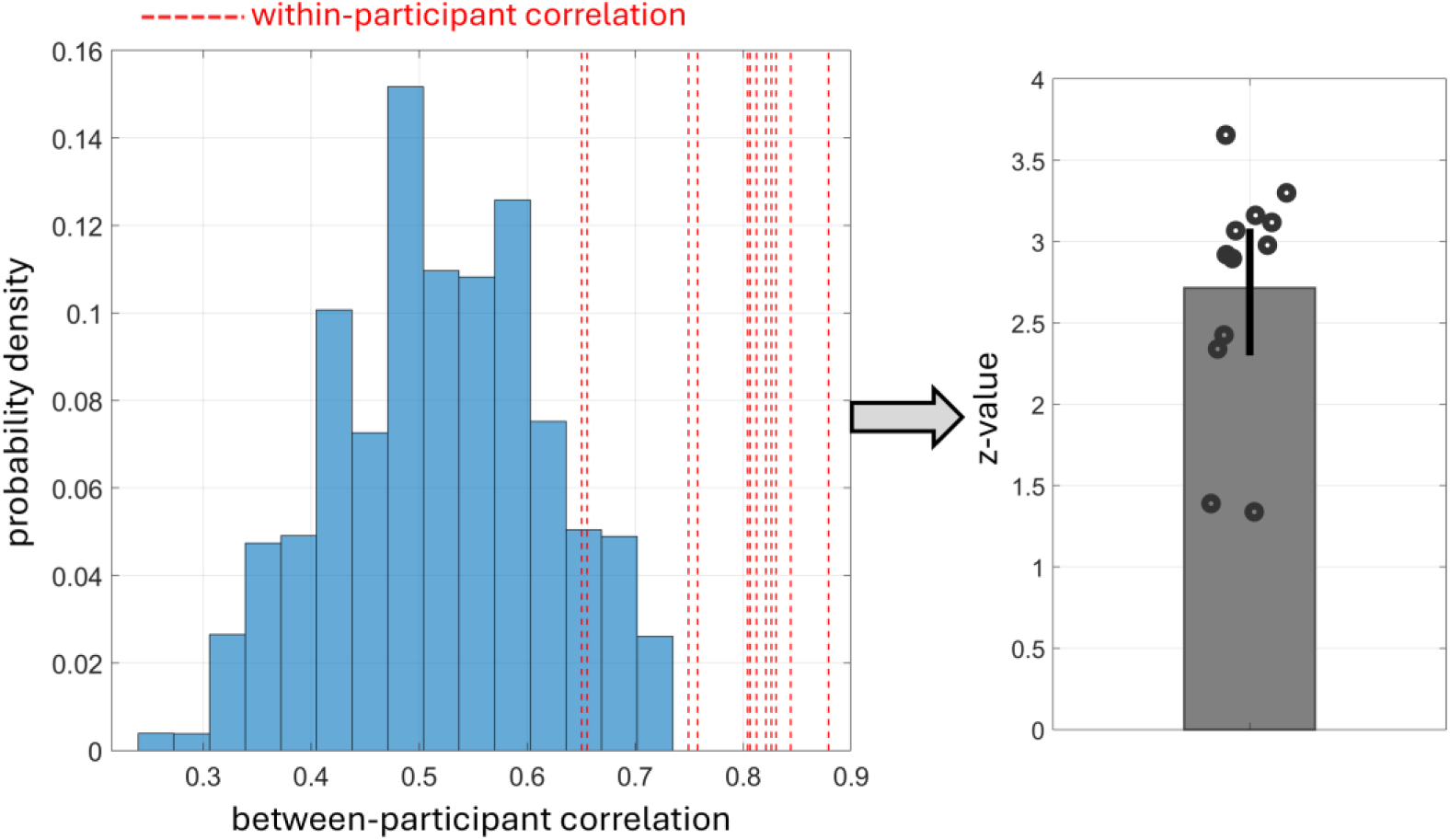
Consistency and uniqueness of the obtained 30×30 perceptual dissimilarity matrices (see Figure 1B) across participants. The left panel shows the histogram of the between-participant correlations computed by randomly correlating the dissimilarity matrix derived from one session of a participant with that of another participant. The dotted red lines indicate the within-participant correlation, representing the correlation between the dissimilarity matrices derived from the first and second sessions of the same participant. The right panel shows the z-values of the within-participant correlations computed relative to the between-participant correlation histogram in the left panel. The bar indicates the mean z-value, the error bar shows its 95% confidence interval, and each dot on the plot corresponds to a participant. The dissimilarity matrices were highly consistent within each individual and unique to them.

Given the self-consistency of similarity judgments, the within-participant correlation could reflect the robustness of the main and alternatively pre-registered methods in estimating the dissimilarity matrix from the collected similarity ranking data. The within-participant correlation was 0.79 ± 0.021 (mean ± SE) when the alternatively pre-registered machine-learning approach was used, whereas it was 0.72 ± 0.018 (mean ± SE) when the main pre-registered approach was used (see Supplementary Figure 5). The within-participant correlation was significantly higher (signed-rank test p-value < 0.001) when the machine-learning approach was used, indicating its superior performance.

Therefore, we presented the results using the dissimilarity matrices derived by the machine-learning approach in the main figures (Figures 2 to 5). The results using the dissimilarity matrices from the main pre-registered approach are instead shown in the supplementary figures (see Supplementary Figures 7 to 10). However, we adhered to our pre-registration and exclusively used the main pre-registered approach at all stages, including selecting the face pairs (see section 2.2.3), examining the data-collection stopping criterion (see section 2.3), and testing our hypotheses. Nevertheless, there were no discrepancies in the results; the stopping criterion was met, and the hypotheses were confirmed using both approaches. The only difference was that the results were stronger with the machine-learning approach, which can be attributed to its more accurate estimation of the dissimilarity matrices as discussed above.

#### 3.1.2. #JNDs

The #JNDs between face pairs were measured once for a participant, so their robustness cannot be evaluated in the same way as the perceptual dissimilarity above. However, given that the #JNDs, unlike the dissimilarity, is considered an objective measure (meaning it is not expected to change with factors like the participant’s state of mind), its robustness can be assessed by the stability of its corresponding staircase, even with a single run. The ratio of reversals within the last 20 trials was about 0.3 ± 0.08 (mean ± SD) across participants in the examined face pairs staircases (see Supplementary Figure 6). Given the 1-up and 2-down protocol used, the theoretical maximum reversal ratio is 0.6, corresponding to the case that each “up” is followed by a “down” and vice versa. While a ratio of 0.6 means convergence within the smallest step size (*S*) (see section 2.2.4), an observed ratio of 0.3 indicates that staircases typically converged within a reasonably tight window on the order of a few *S* (roughly 3 × *S*). As a reference, the example staircase shown in Figure 1C has a reversal ratio of 0.35 within its last 20 trials. Taken together, these suggest that the staircases converged fairly well across participants. Therefore, the measured #JNDs were reasonably robust.

### 3.2. The relationship between perceptual discrimination capacity and subjective perceptual dissimilarity

We found a strong positive association between individuals’ perceptual discrimination capacity and their subjective perceptual dissimilarity (Figure 3). Face pairs that were harder to discern in the near-threshold task were subjectively perceived as more similar in the (supra-threshold) similarity judgment task. This association was highly significant at the group level. The mean z-value was 2.63, with a 95% confidence interval of [2.18, 3.10]. Other complementary statistical tests also indicated high significance: a t-test on the z-values yielded a p-value < 0.00001 and a BF > 10000 (one-tailed; assuming the opposite side is theoretically implausible), and a Fisher’s test combining individual-level p-values (one-tailed) resulted in a p-value < 0.00001. Significance was achieved at the individual level (p-value < 0.05, one-tailed) in most (eleven) participants. This corresponds to a Bayesian population prevalence of 0.91 with a 95% highest posterior density interval of [0.66, 0.99], indicating that the true likelihood of observing an individual-level significance, following our experimental design, lies between 66% and 99%.

**Figure 3.**
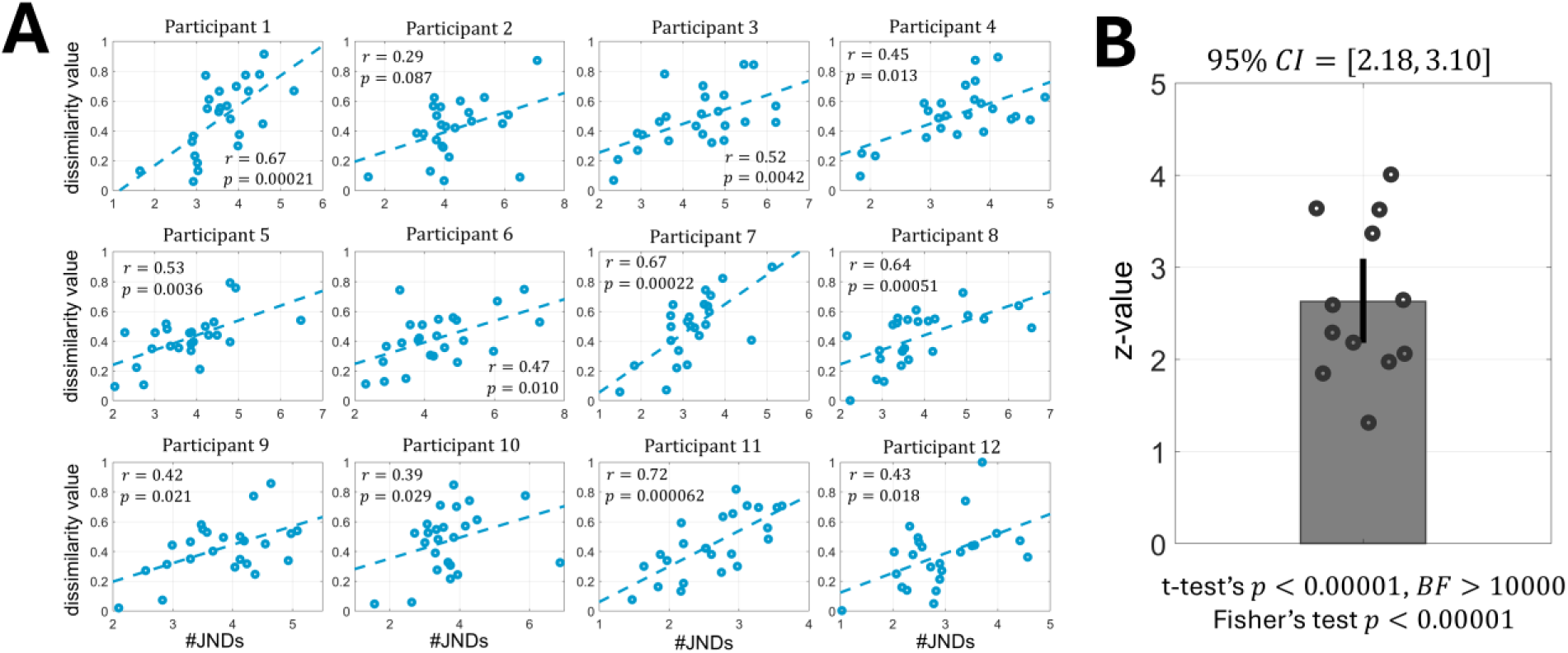
Relationship between perceptual discrimination capacity and subjective perceptual dissimilarity in each participant. **(A)** Each subplot illustrates the relation between perceptual discrimination capacity, quantified in #JNDs (x-axis), and subjective perceptual dissimilarity (y-axis) in each participant, with each data point corresponding to a face pair. r indicates the Spearman correlation coefficient, and p denotes its associated p-value (one-tailed, assuming negative correlations are theoretically implausible). **(B)** Individuals’ Spearman correlations in A were transformed to z-values for group-level hypothesis testing. The bar shows the mean z-value across participants, the error bar represents the 95% confidence interval of this group-mean z-value, calculated through bootstrapping, and each dot on the plot corresponds to a participant. As displayed under the plot, analysis of the group-level effect by applying a t-test on the z-values yielded a p-value smaller than 0.00001 and a Bayes factor (BF) above 10000 (one-tailed). Additionally, employing a Fisher’s combined probability test, combining the individual-level p-values, similarly resulted in a p-value smaller than 0.00001. The results suggest a strong positive association between perceptual discrimination capacity and subjective perceptual dissimilarity, confirming Hypothesis 1.

Importantly, we found that the association between perceptual discrimination capacity and subjective perceptual dissimilarity was individual-specific. As depicted in Figure 4, one’s subjective perceptual dissimilarity was better correlated with one’s own perceptual discrimination capacity than with that of others. This finding was significant at the group level, with a group-mean z-value of 1.19 and a 95% confidence interval of [0.77, 1.59]. Additional complementary statistics further highlighted the high significance of the result: a t-test on the z-values resulted in a p-value of 0.00010 and a BF of 306 (one-tailed; the opposite side is theoretically meaningless), and a Fisher’s combined probability test yielded a p-value of 0.000062. At the individual-level, this finding reached significance (p-value < 0.05, one-tailed) in five participants, equating to a Bayesian population prevalence highest posterior density interval of [0.14, 0.66]. This implies that, given our experimental design and factors such as measurement noise, the true likelihood of observing an individual-level significance lies between 14% and 66%. Note that although only five (out of twelve) participants reached the individual-level significance threshold, most remaining participants showed the trend. Accordingly, the group-level statistic was highly significant, supporting our Hypothesis 2.

**Figure 4.**
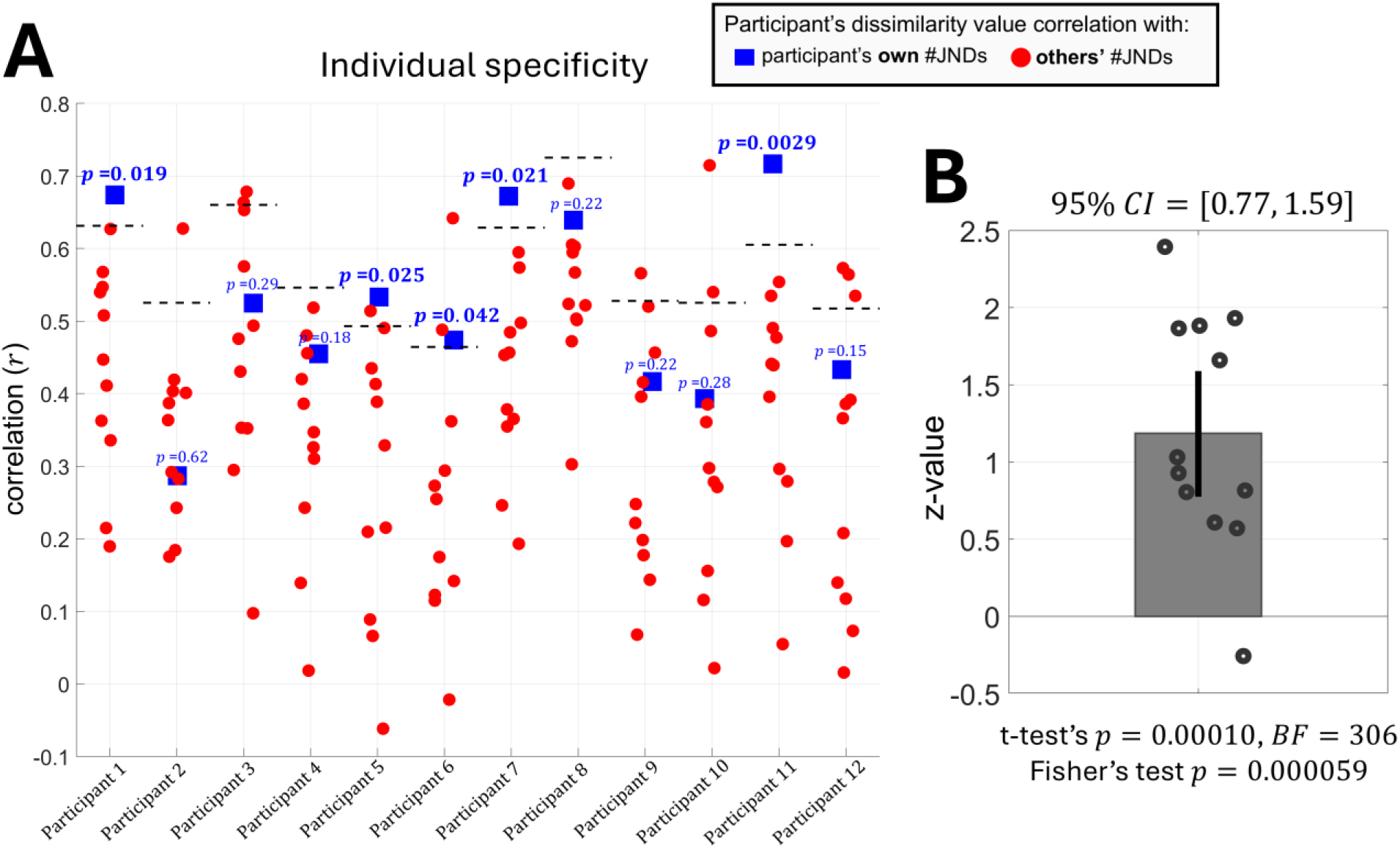
Individual-specificity of the relationship between perceptual discrimination capacity and subjective perceptual dissimilarity. **(A)** The blue square and red circle indicate the Spearman correlation coefficient between the dissimilarity value in each participant and their own #JNDs and other participants’ #JNDs, respectively. The dotted horizontal black line denotes the Spearman correlation value at which a permutation test yielded a p-value of 0.05, rejecting a null hypothesis of no individual-specificity. The null hypothesis distribution consisted of possible Spearman correlations between the dissimilarity value in the target participant and others’ #JNDs. The p-value rejecting the null hypothesis in each participant is shown in blue at the top of the blue squares (one-tailed; as the opposite side, where others’ #JNDs better predict an individual’s dissimilarity, is theoretically meaningless). **(B)** Individuals’ specificity statistics in A were converted to z-values for group-level hypothesis testing. The bar shows the group-mean z-value, the error bar indicates its 95% confidence interval, and each dot corresponds to a participant. The confidence interval was above zero, indicating group-level significance. This demonstrates that the association between perceptual discrimination capacity and subjective perceptual dissimilarity is quite specific to each individual, confirming Hypothesis 2. The Other complementary statistics, similar to Figure 3B, are displayed below the bar plot, further highlighting the strong group-level significance.

Figure 5 illustrates the association between perceptual discrimination capacity and subjective perceptual dissimilarity in each face pair across participants. This association was positive in most pairs and reached a p-value < 0.05 (one-tailed; assuming only positive associations are theoretically meaningful) in seven pairs. A t-test on the positivity of the association across pairs yielded a p-value of 0.00010 and a BF of 279 (one-tailed). These observations further support our hypotheses, indicating that individuals with a lower capacity for discriminating a pair tended to judge that pair as more similar.

**Figure 5.**
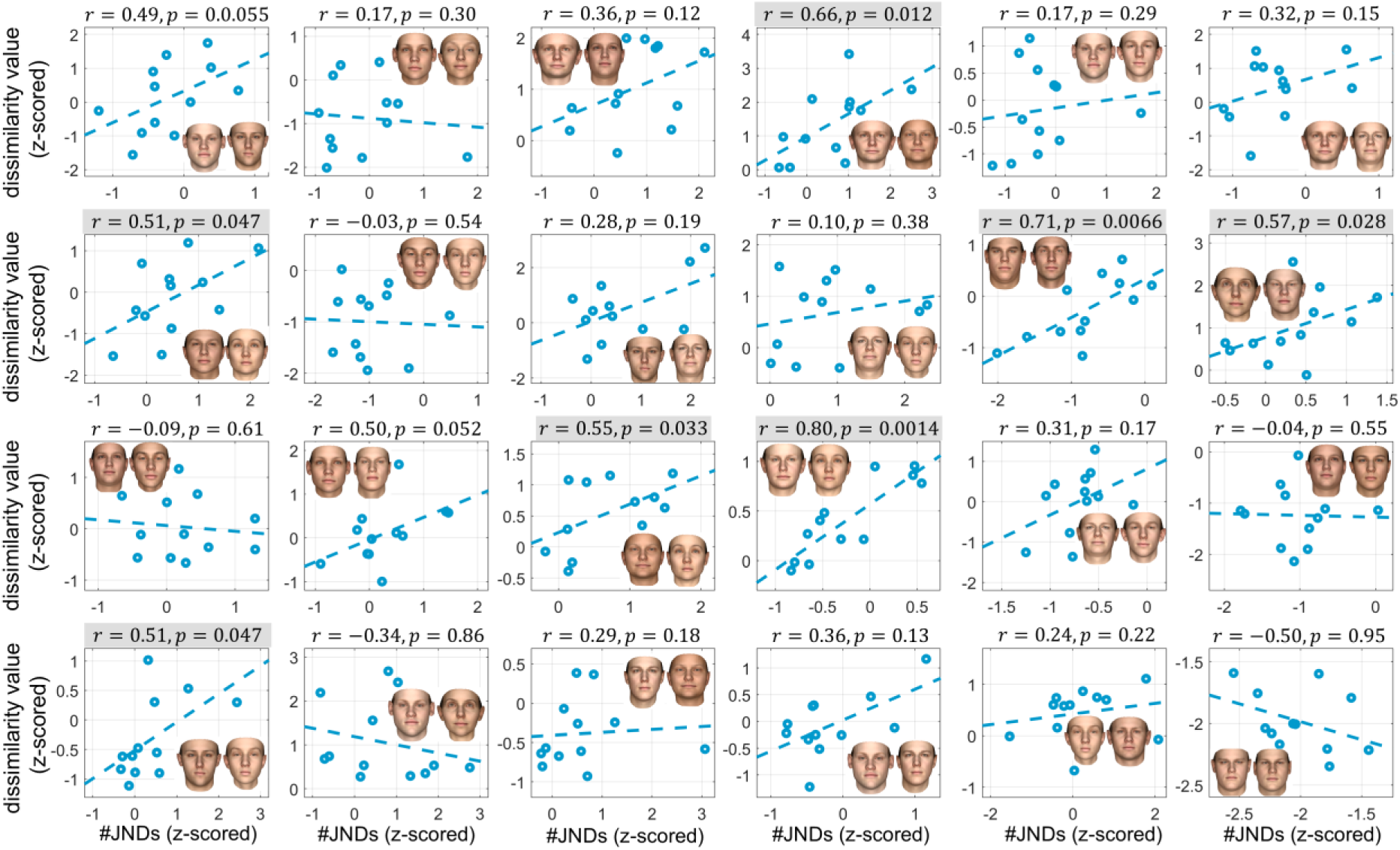
Relationship between perceptual discrimination capacity and subjective perceptual dissimilarity across participants in each face pair. Each subplot represents the association between the #JNDs and dissimilarity value across participants in a given face pair, with each dot corresponding to a participant. The dissimilarity value and #JNDs were z-scored within each participant to ensure a consistent scaling across participants. r and p displayed on top of each subplot indicate the Spearman correlation coefficient and its associated p-value (one-tailed, assuming only positive correlations are theoretically meaningful), respectively. Pairs with a p-value < 0.05 are highlighted in gray. The subplots are organized such that the face pair with the most controversial dissimilarity value across participants is shown in the top left, and the pair with the least controversial value is shown in the bottom right. A positive association between perceptual discrimination capacity and subjective perceptual dissimilarity was observed in most pairs, indicating that participants with a lower capacity for discriminating a pair tended to judge that pair as more similar, which supports our hypotheses. A t-test on the r value being positive across pairs yielded a p-value of 0.00010 and a BF of 279 (one-tailed). We note that this positive association was more salient in the more controversial pairs (e.g., compare the first three rows with the last row).

The association was more apparent in the controversial pairs (i.e., those with high-variance perceptual dissimilarity across participants) than in the less-controversial pairs. As shown in Figure 5, the weakest correlations were observed in the less-controversial pairs (see the last row in this figure). Specifically, for the controversial pairs (first three rows), the mean correlation was 0.35 ± 0.063 [SE], whereas for the less-controversial pairs (last row), it was 0.094 ± 0.17 [SE]. A two-sample t-test indicated a trend-level difference (p = 0.09, BF = 1.2), with higher correlation values for the controversial pairs.

This outcome was expected because psychophysical measures (i.e., #JNDs and perceptual dissimilarity) inevitably contain noise, which can obscure true relationships that may exist, particularly when variability is small across participants. Despite this intuitive expectation, six less-controversial pairs were intentionally examined in the study to assess whether the hypothesized relationship could still be observed in them, given the measurement noise. The results suggest that while our current design is sufficiently sensitive to detect the individual-specific association between perceptual discrimination capacity and subjective perceptual dissimilarity in controversial pairs, it may lack the resolution when the pairs are less-controversial, assuming our hypothesis holds across all pairs. We measured #JNDs once, whereas perceptual dissimilarity was measured twice across two sessions and subsequently averaged. Therefore, conducting further repeated measures, especially for #JNDs, seems necessary to potentially capture the association across all pairs.

Finally, to remain completely faithful to our pre-registered plan, we also tested using a 2D MDS (rather than a 5D MDS) to estimate dissimilarity values. We hoped that a 2D MDS, by representing the dissimilarity relations in only two dimensions, would preserve the most robust aspects of these relations, thereby making their association with perceptual discrimination capacity more salient. However, as shown in Supplementary Figure 11, this was not the case. Although the association, as well as its individual specificity, were significant, they became weaker when a 2D MDS was used. Not only that, the within-participant correlation of the dissimilarity matrices, along with their individual uniqueness (evaluated by comparison to the between-participant correlation), also weakened. Therefore, these results suggest that further reducing dimensionality to two dimensions caused more harm (information loss) than benefit (e.g., noise reduction).

## 4. Discussion

In the present study, we used a near-threshold psychophysical task to quantify perceptual discrimination capacity, which indicates one’s capability to distinguish two stimuli (Figure 1C). We examined whether this perceptual discrimination capacity measured at near-threshold is associated with subjective perceptual similarity rankings (Figure 1B) given at suprathreshold (Hypothesis 1). More critically, we explored whether this association is specific to each individual, meaning that one’s subjective perceptual similarity judgements are better explained by their own perceptual discrimination capacities than by those of others (Hypothesis 2). We first conducted a small pilot study, which provided initial support for both hypotheses (see Supplementary Figure 3). Then, the results from our main study confirmed them (Hypothesis 1 in Figure 3 and Hypothesis 2 in Figure 4).

Perceptual capacities were measured via a task with objective responses, whereas perceptual similarity judgments were measured in a task with a free nature and no defined correct answers. Therefore, our results suggest that perceptual capacities may serve as a quasi-objective ground-truth basis for similarity judgments. Consequently, although subjective, these judgments could be labeled as more ‘accurate’ if they are better aligned with one’s own perceptual capacities.

Perceptual discrimination capacity reflects an individual’s sensory limits and is therefore constrained by the underlying neuronal tunings in the visual system (in our context of visual stimuli). Similarity judgment, on the other hand, involves a cognitive evaluation of stimuli. Accordingly, perceptual capacity can be viewed as a marker of visual system resolution, while similarity judgments represent a higher-level decision informed by those markers. In this sense, our findings may suggest that similarity judgments are somewhat ‘metacognitive’ – in a limited sense – reflecting *implicit* readouts of one’s own perceptual capacities. Relatedly, instability (noise) in perceptual similarity judgments could be considered as the result of inaccurate self-assessment of one’s own perceptual capacities.

Consequently, higher cortical brain areas, particularly the prefrontal cortex, may play a critical role in perceptual similarity judgments, given that its activity has been demonstrated to be associated with perceptual metacognition (Fleming et al., 2014; McCurdy et al., 2013; Morales et al., 2018). Of course, the current study does not directly test this hypothesis about neural mechanisms. Others have suggested that perceptual similarity information resides within the sensory cortices (Malach, 2021). In light of this, we are currently investigating whether perceptual similarity representations can be found beyond the visual areas, such as the lateral prefrontal cortex, using fMRI.

Our results could also shed light on one conundrum regarding large language models and consciousness. Recently, it has been reported that models built with current artificial intelligence technology can produce human-like similarity ratings (Kawakita et al., 2023; Marjieh et al., 2023). If the qualitative characters of conscious perception are determined by the relevant similarity relations, as some researchers assume (Clark, 2000; Lau et al., 2022; Malach, 2021; Moharramipour & Lau, 2024; Rosenthal, 2010; Tallon-Baudry, 2022; Zeleznikow-Johnston et al., 2023), does it mean that these artificial agents are conscious (Moharramipour & Lau, 2024)? Or, at least, does it mean that they contain the essential information that is encapsulated within human perceptual experiences? The answer is probably no, considering the perspective described above. That is, for similarity judgments to be relevant for subjective experiences, according to our findings, they need to reflect one’s own perceptual capacities. What these models do is simply to mimic what humans say in general, and as such, their similarity judgments at best reflect common world knowledge about the physical characteristics of the stimuli, but they are not about one’s own perceptual capacities (of which these models have none). There is, thus, a critical difference between humans and those models, in terms of what the similarity judgments mean for them.

From a structural perspective, similarity relations are commonly represented in a metric psychological space (Borg & Groenen, 2005; Hebart et al., 2020; Schurgin et al., 2020). Our results suggest that for each individual, the warping and scale of this space are tied to their perceptual capacity. Furthermore, as formulated by Roger Shepard, generalization decays exponentially with increasing distance in psychological space (Shepard, 1987). In this framework, higher perceptual capacity, by increasing the psychological distance, could cause a stronger reduction in generalization (a steeper exponential decay).

Finally, we note that we used face stimuli in our study because they are highly complex (i.e., high-dimensional) and commonly encountered in daily life. However, it may be necessary to examine how our findings hold in other types of visual stimuli, such as colors or artificially constructed stimuli (e.g., Greebles (Gauthier & Tarr, 1997)), and even across different sensory domains (Faivre et al., 2018). Moreover, future studies could use multi-featural stimuli that, unlike faces, do not generally rely on holistic perception, such as stimuli with varying shapes, sizes, and colors, to assess how perceptual capacity within each individual feature influences similarity judgment.

## Author contributions

A.M. contributed to conceptualization, project planning and design, methodology application, data collection, data analysis, visualization, writing, review, and editing.

W.Z. contributed to methodology application, data analysis, review, and editing.

D.R. contributed to conceptualization, project design, review, and editing.

H.L. contributed to conceptualization, supervision, project planning, design, analysis, review, and editing.

## Data and code availability

The data and the code used in the study are publicly accessible from the GitHub repository below: https://github.com/AliMoharramipour/Subjective-Dissimilarity-and-Discrimination-Capacity-

## Supplementary Materials

**Supplementary Table 1.**
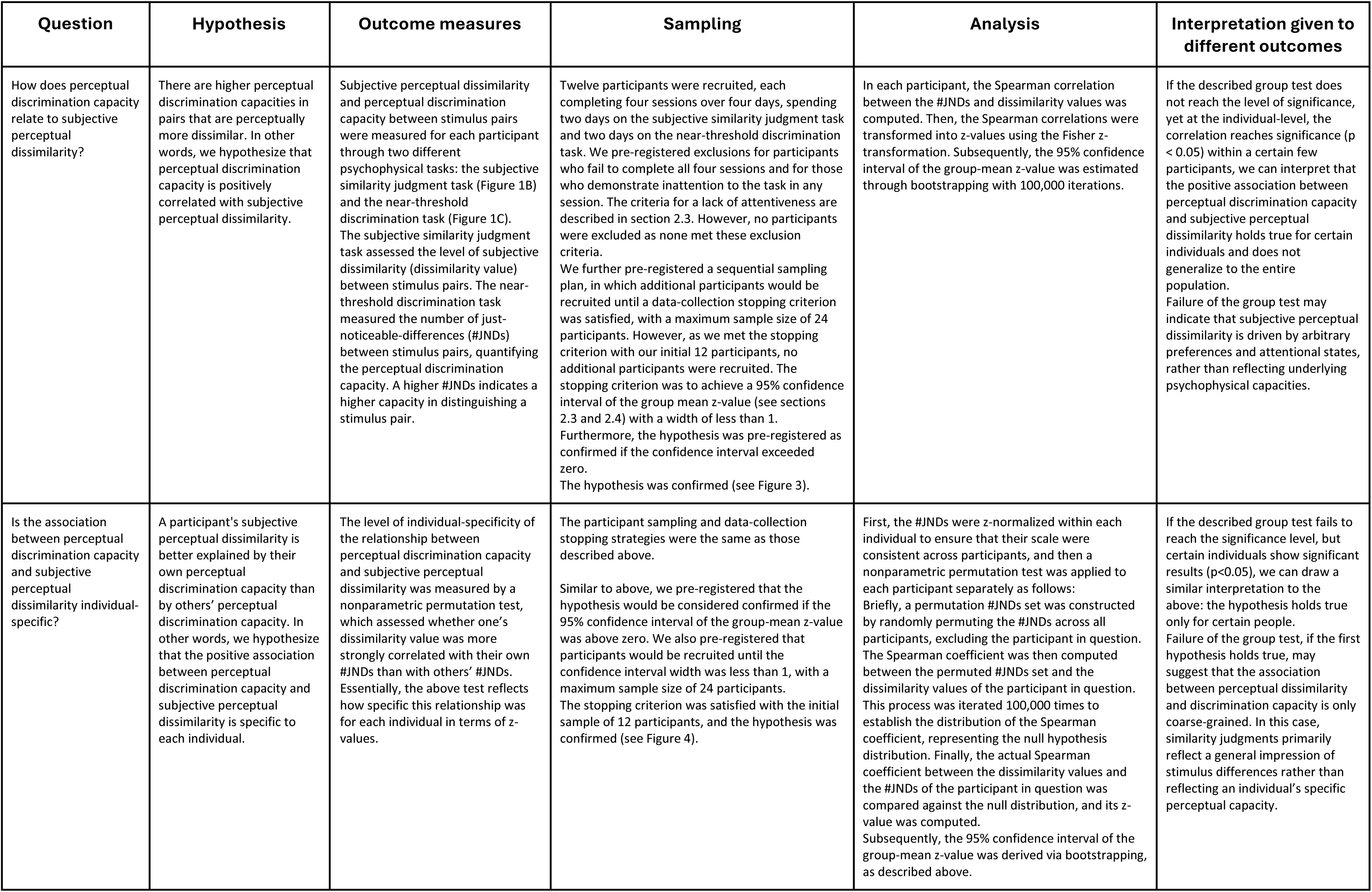
Experimental Design Table.

| Question | Hypothesis | Outcome measures | Sampling | Analysis | Interpretation given to different outcomes |
| --- | --- | --- | --- | --- | --- |
| How does perceptual discrimination capacity relate to subjective perceptual dissimilarity? | There are higher perceptual discrimination capacities in pairs that are perceptually more dissimilar. In other words, we hypothesize that perceptual discrimination capacity is positively correlated with subjective perceptual dissimilarity. | Subjective perceptual dissimilarity and perceptual discrimination capacity between stimulus pairs were measured for each participant through two different psychophysical tasks: the subjective similarity judgment task (Figure 1B) and the near-threshold discrimination task (Figure 1C). The subjective similarity judgment task assessed the level of subjective dissimilarity (dissimilarity value) between stimulus pairs. The near-threshold discrimination task measured the number of just-noticeable-differences (#JNDs) between stimulus pairs, quantifying the perceptual discrimination capacity. A higher #JNDs indicates a higher capacity in distinguishing a stimulus pair. | Twelve participants were recruited, each completing four sessions over four days, spending two days on the subjective similarity judgment task and two days on the near-threshold discrimination task. We pre-registered exclusions for participants who fail to complete all four sessions and for those who demonstrate inattention to the task in any session. The criteria for a lack of attentiveness are described in section 2.3. However, no participants were excluded as none met these exclusion criteria.<br>We further pre-registered a sequential sampling plan, in which additional participants would be recruited until a data-collection stopping criterion was satisfied, with a maximum sample size of 24 participants. However, as we met the stopping criterion with our initial 12 participants, no additional participants were recruited. The stopping criterion was to achieve a 95% confidence interval of the group mean z-value (see sections 2.3 and 2.4) with a width of less than 1. Furthermore, the hypothesis was pre-registered as confirmed if the confidence interval exceeded zero.<br>The hypothesis was confirmed (see Figure 3). | In each participant, the Spearman correlation between the #JNDs and dissimilarity values was computed. Then, the Spearman correlations were transformed into z-values using the Fisher z-transformation. Subsequently, the 95% confidence interval of the group-mean z-value was estimated through bootstrapping with 100,000 iterations. | If the described group test does not reach the level of significance, yet at the individual-level, the correlation reaches significance ($p < 0.05$) within a certain few participants, we can interpret that the positive association between perceptual discrimination capacity and subjective perceptual dissimilarity holds true for certain individuals and does not generalize to the entire population.<br>Failure of the group test may indicate that subjective perceptual dissimilarity is driven by arbitrary preferences and attentional states, rather than reflecting underlying psychophysical capacities. |
| Is the association between perceptual discrimination capacity and subjective perceptual dissimilarity individual-specific? | A participant's subjective perceptual dissimilarity is better explained by their own perceptual discrimination capacity than by others' perceptual discrimination capacity. In other words, we hypothesize that the positive association between perceptual discrimination capacity and subjective perceptual dissimilarity is specific to each individual. | The level of individual-specificity of the relationship between perceptual discrimination capacity and subjective perceptual dissimilarity was measured by a nonparametric permutation test, which assessed whether one's dissimilarity value was more strongly correlated with their own #JNDs than with others' #JNDs. Essentially, the above test reflects how specific this relationship was for each individual in terms of z-values. | The participant sampling and data-collection stopping strategies were the same as those described above.<br>Similar to above, we pre-registered that the hypothesis would be considered confirmed if the 95% confidence interval of the group-mean z-value was above zero. We also pre-registered that participants would be recruited until the confidence interval width was less than 1, with a maximum sample size of 24 participants. The stopping criterion was satisfied with the initial sample of 12 participants, and the hypothesis was confirmed (see Figure 4). | First, the #JNDs were z-normalized within each individual to ensure that their scale were consistent across participants, and then a nonparametric permutation test was applied to each participant separately as follows: Briefly, a permutation #JNDs set was constructed by randomly permuting the #JNDs across all participants, excluding the participant in question. The Spearman coefficient was then computed between the permuted #JNDs set and the dissimilarity values of the participant in question. This process was iterated 100,000 times to establish the distribution of the Spearman coefficient, representing the null hypothesis distribution. Finally, the actual Spearman coefficient between the dissimilarity values and the #JNDs of the participant in question was compared against the null distribution, and its z-value was computed.<br>Subsequently, the 95% confidence interval of the group-mean z-value was derived via bootstrapping, as described above. | If the described group test fails to reach the significance level, but certain individuals show significant results ($p < 0.05$), we can draw a similar interpretation to the above: the hypothesis holds true only for certain people.<br>Failure of the group test, if the first hypothesis holds true, may suggest that the association between perceptual dissimilarity and discrimination capacity is only coarse-grained. In this case, similarity judgments primarily reflect a general impression of stimulus differences rather than reflecting an individual's specific perceptual capacity. |

**Supplementary Figure 1.**
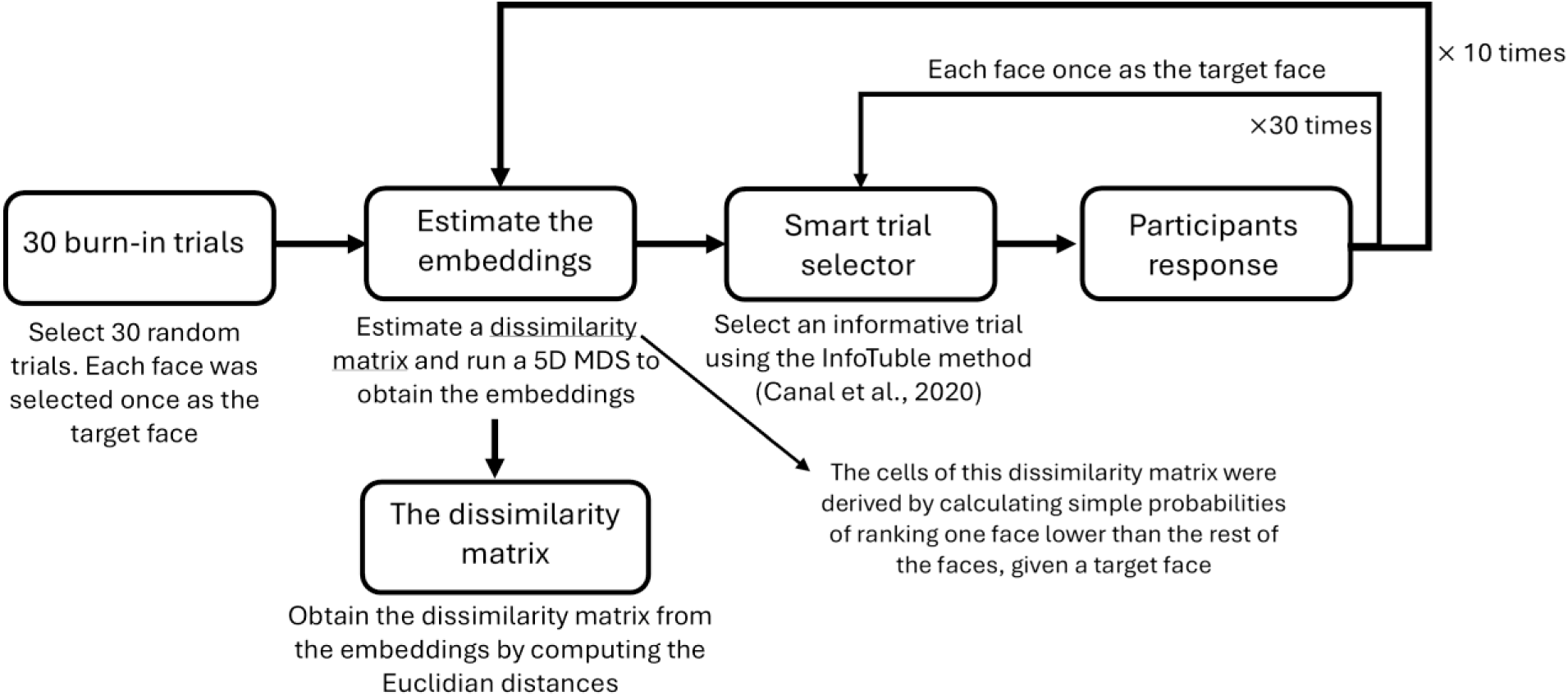
Schematic of the subjective similarity judgment task design.

**Supplementary Figure 2.**
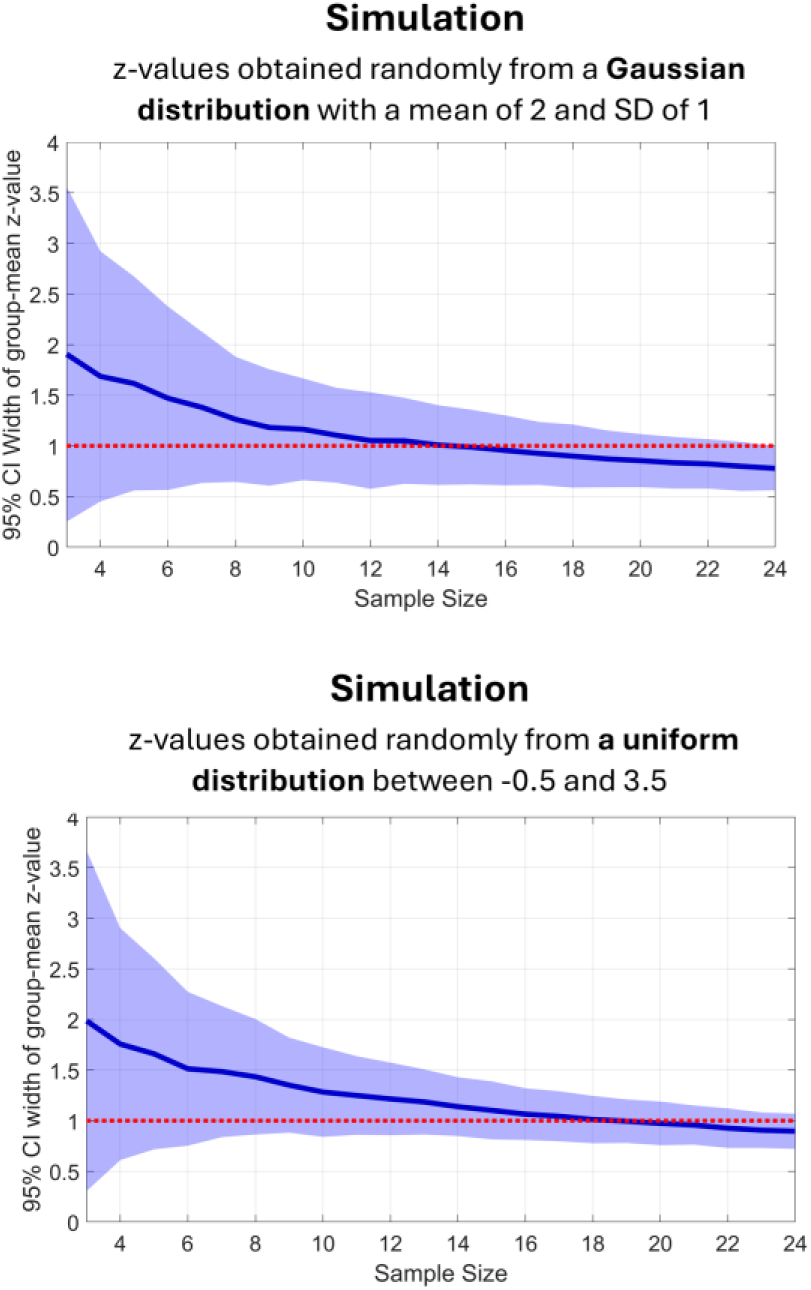
Simulations to determine a feasible sample size for our data collection stopping criterion. We ran two simulations: one generated z-values from a Gaussian distribution with a mean of 2 and SD of 1, and another from a uniform distribution ranging between -0.5 and 3.5. These example distributions appeared reasonable given our expectations from the pilot data and were conservative enough. For example, the 95% confidence interval (CI) width of the group-mean z-value in our pilot data with four participants was around 1.5; however, in the presented simulations, the 95% CI width is, on average, around 1.7 for the same sample size of four. The shaded area indicates the 2.5th and 97.5th percentiles of the 95% CI width obtained over 1000 simulations. Assuming the used distributions are realistic, there was a high likelihood of hitting the stopping criterion by reaching a maximum sample size of 24. We, in fact, reached the stopping criterion (i.e., a CI width smaller than 1) with our initial 12 participants in the main study.

## Pilot data

We conducted a pilot study with four participants prior to pre-registration (i.e., publishing the Stage 1 registered report). The experimental design was similar to the main experiment (Figure 1), with a few differences. Participants performed the subjective similarity judgment task only once. After all of them completed the task, 13 pairs of faces were selected, and the participants were all invited back to perform, in a different session, the near-threshold discrimination task on these pairs. In this pilot study, we randomly selected the pairs while ensuring that most of them have high dissimilarity values SD across participants, indicating that the degree of subjective dissimilarity is quite ‘controversial’. The subjective similarity judgment task and the near-threshold discrimination task were conducted similarly to the main experiment, except that there was no time constraint on both tasks, and no trial-by-trial behavioral feedback was provided during the near-threshold discrimination task. Additionally, for one participant, the near-threshold discrimination task comprised 50 rather than 60 trials. We applied the same analysis approach described in section 2.4 to the pilot data.

Hypothesis 1 was confirmed in the pilot study (Supplementary Figures 3A and 3B), indicating that subjective perceptual dissimilarity and perceptual discrimination capacity are highly correlated. The correlation was significant (p-value < 0.05; one-tailed) at the individual-level in 3 out of 4 participants. At the group-level, the mean z-value across participants was 2.33, with a 95% confidence interval between 1.65 and 3.02. A t-test on the z-values yielded a p-value of 0.0062, and a Bayes factor (BF) of 12 (one-tailed). Moreover, a Fisher’s test, combining the individual-level p-values, resulted in a p-value of 0.000011.

More importantly, Hypothesis 2 was also confirmed in the pilot study (Supplementary Figures 3C and 3D), suggesting that the association between subjective perceptual dissimilarity and perceptual discrimination capacity is specific to each individual. The statistic was significant (p-value < 0.05; one-tailed) at the individual-level in 3 out of 4 participants. At the group-level, the mean z-value across participants was 2.26, with a 95% confidence interval between 1.46 and 2.94. A t-test on the z-values resulted in a p-value of 0.0081 and a BF of 10 (one-tailed), and a Fisher’s test yielded a p-value smaller than 0.00001.

**Supplementary Figure 3.**
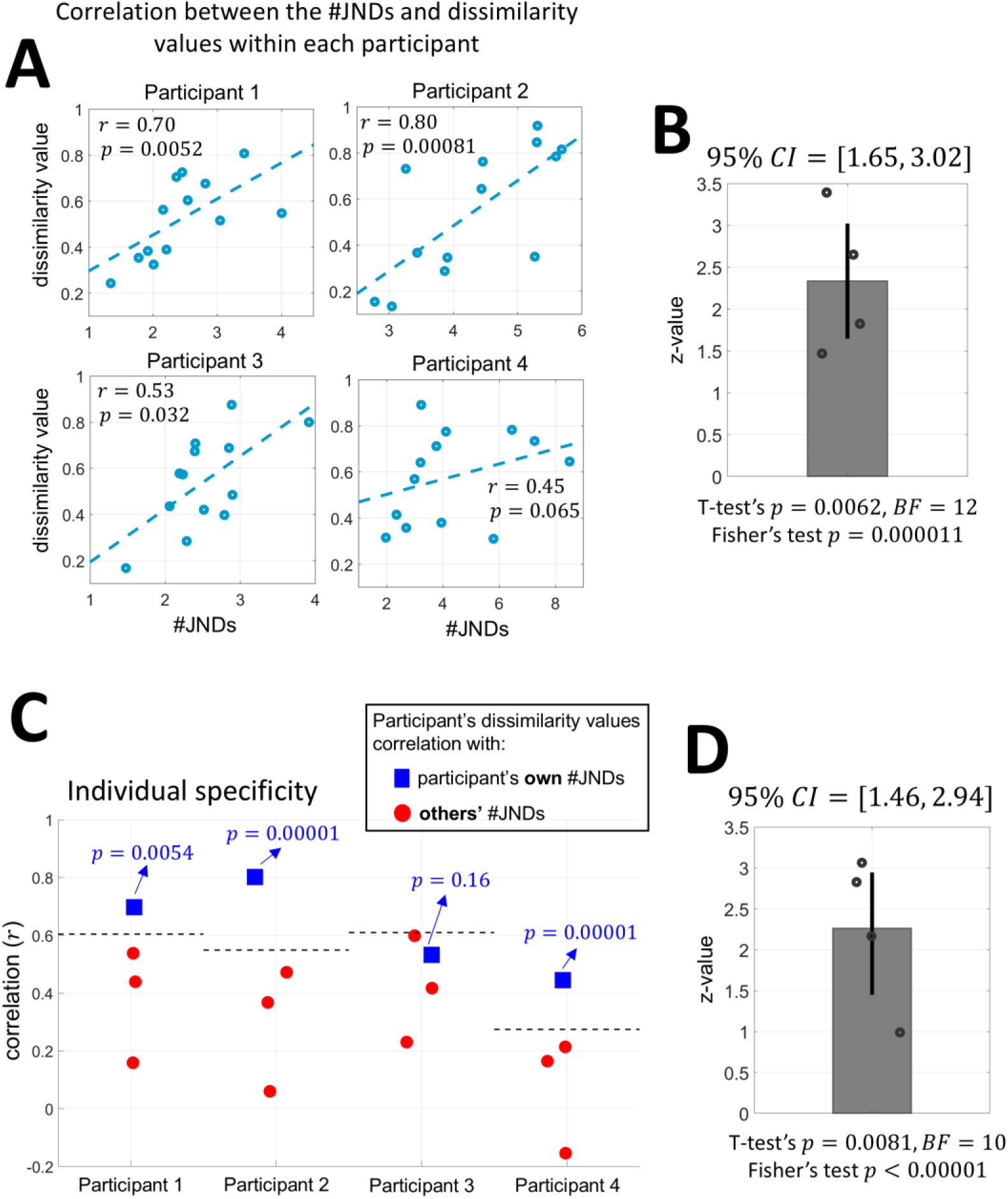
Results in the pilot study with N = 4 participants, who performed the subjective similarity judgment task once in a session and then the near-threshold discrimination task once in another session on 13 pre-selected pairs. **(A,B)** Relationship between perceptual discrimination capacity and subjective perceptual dissimilarity in each participant. The descriptions of the plots are similar to those in Figure 3. **(C,D)** Individual specificity of the relationship between perceptual discrimination capacity and subjective perceptual dissimilarity. The remaining descriptions of the plots are similar to those in Figure 4.

We further explored the correlation between subjective perceptual dissimilarity and perceptual discrimination capacity in each face pair across participants in our pilot study (Supplementary Figure 4). Given the small sample size (i.e., four participants), no meaningful statistical conclusions could be inferred. However, it is notable that the correlations were strongly positive, particularly for the controversial pairs: those with high variability in subjective dissimilarity across participants. Essentially, this also indicated that one’s perceptual discrimination capacity can account for their subjective perceptual dissimilarity: similar inter-individual differences observable in subjective perceptual dissimilarity could also be found in perceptual discrimination capacity.

**Supplementary Figure 4.**
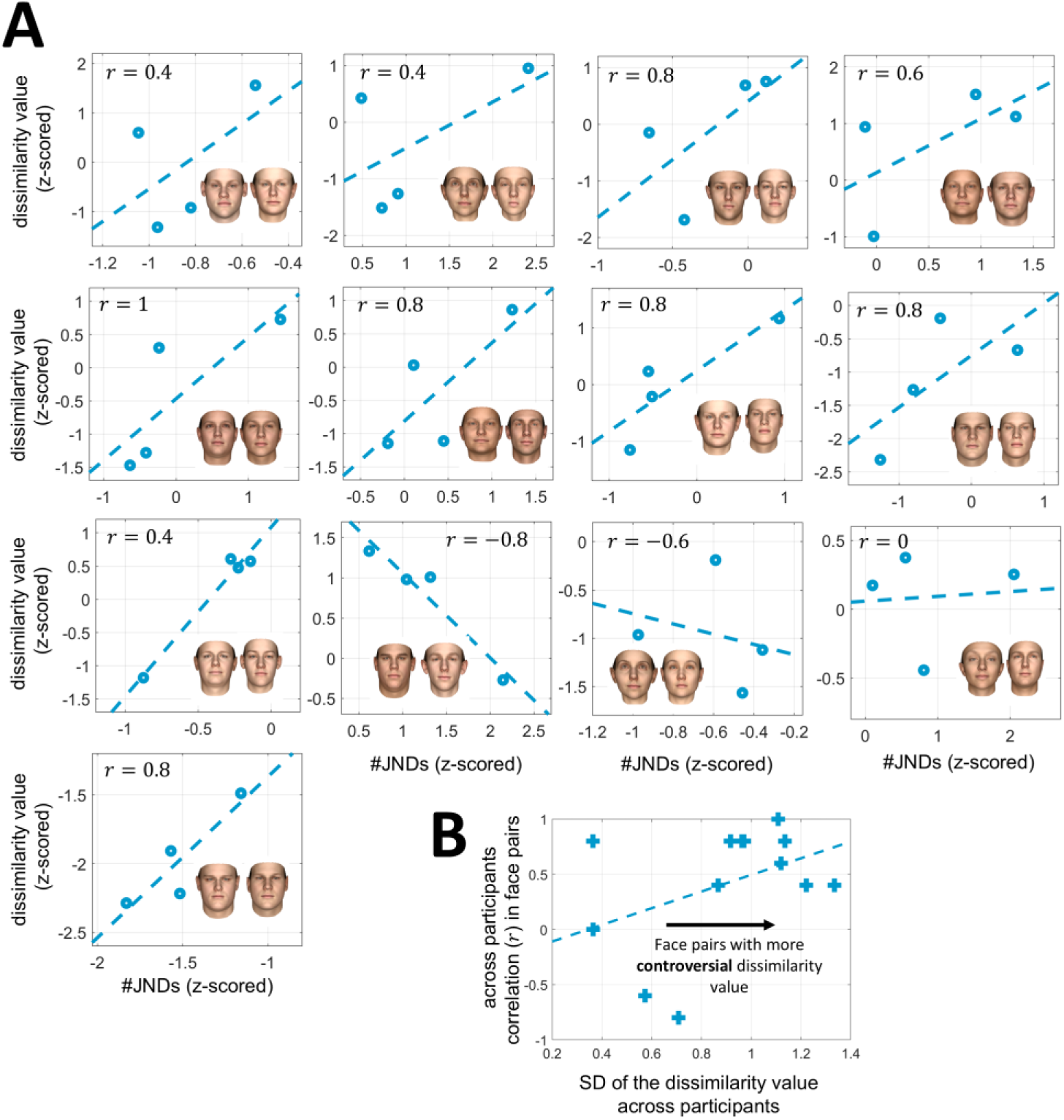
Relationship between perceptual discrimination capacity and subjective perceptual dissimilarity across participants in each face pair (pilot study, N = 4 participants). **(A)** The descriptions of the plots are similar to those in Figure 5. **(B)** Relationship between the face pairs Spearman correlation coefficient shown in A and their controversy in subjective dissimilarity across participants (i.e., SD of dissimilarity values across participants). The association between perceptual discrimination capacity and subjective perceptual dissimilarity across participants was more salient in highly controversial pairs.

**Supplementary Figure 5.**
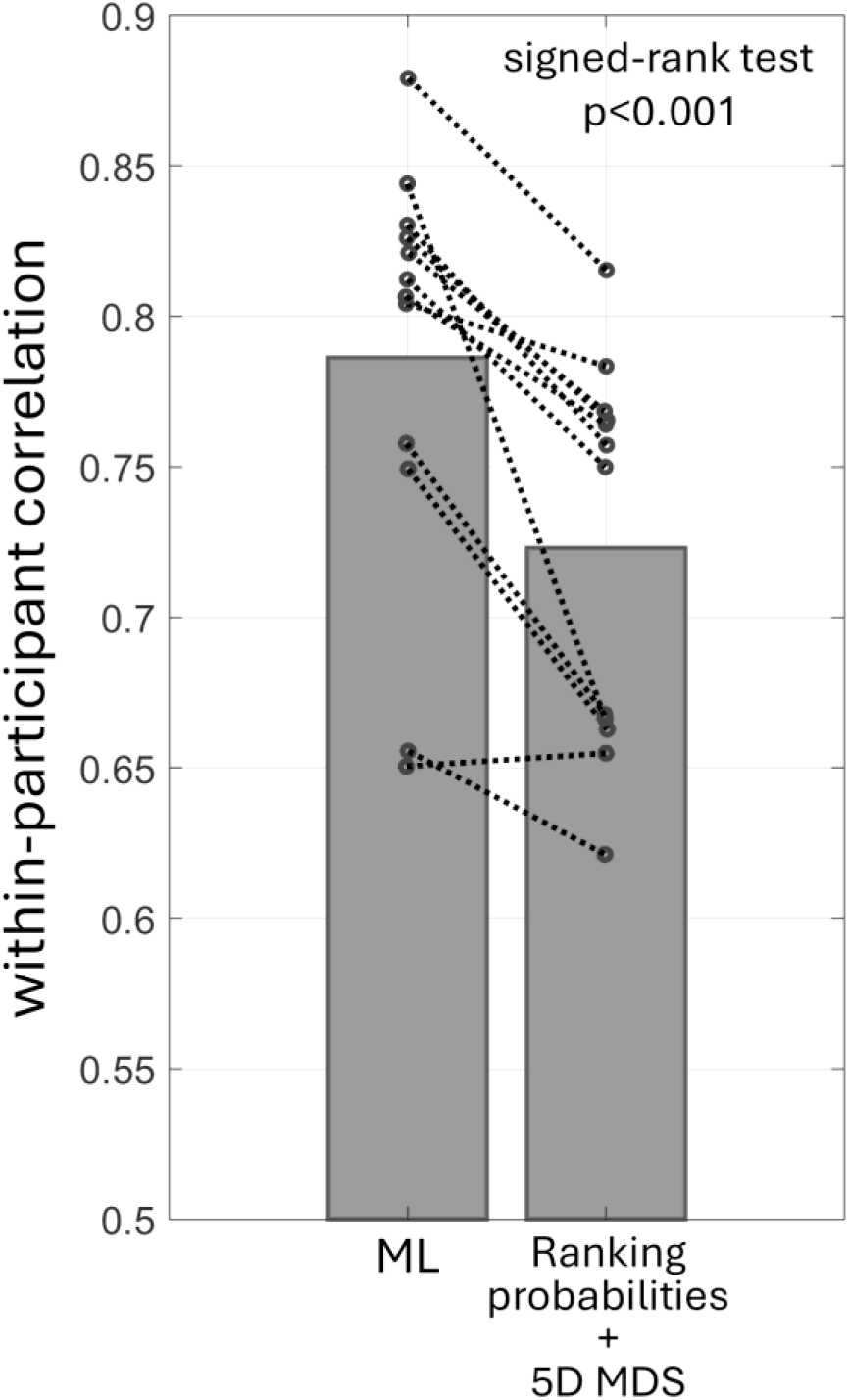
Comparison of the alternatively pre-registered machine-learning approach and the main pre-registered approach in estimating the dissimilarity matrix. The correlation between the dissimilarity matrices from the first and second sessions of the same participants (i.e., within-participant correlation) was higher when derived using the machine-learning approach (signed-rank test p-value < 0.001). Thus, the machine-learning approach could more robustly estimate the dissimilarity matrix from the collected subjective similarity ranking data. To recap, the main pre-registered approach involved estimating ranking probabilities and then applying a 5D MDS to reduce dimensionality and thereby reduce noise. In contrast, the alternatively pre-registered machine-learning approach was designed to directly find the underlying embeddings by minimizing a loss function that penalizes mismatches between the similarity rankings inferred from the embeddings and those made by the participant.

**Supplementary Figure 6.**
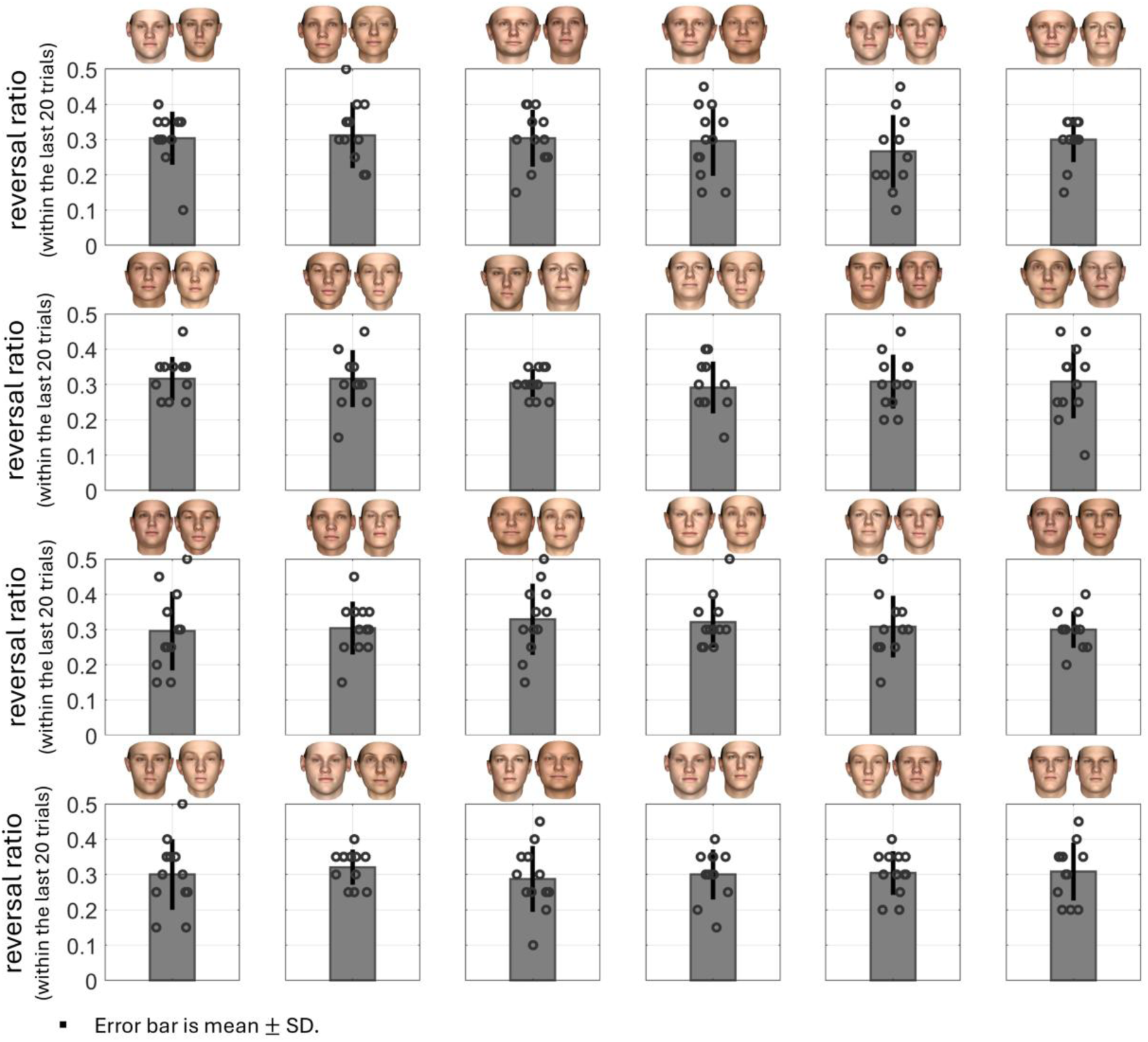
Quality of the measured #JNDs (stability of the staircases). Each panel displays the ratio of reversals within the last 20 trials of each face pair’s staircase. The bar shows the mean value, and the error bar displays its standard deviations (mean ± SD) across participants, with each dot representing a participant. Note that the maximum possible theoretical ratio is 0.6, given the 1-up and 2-down protocol used. As a reference, the example staircase shown in Figure 1C has a reversal ratio of 0.35 within its last 20 trials.

**Supplementary Figure 7.**
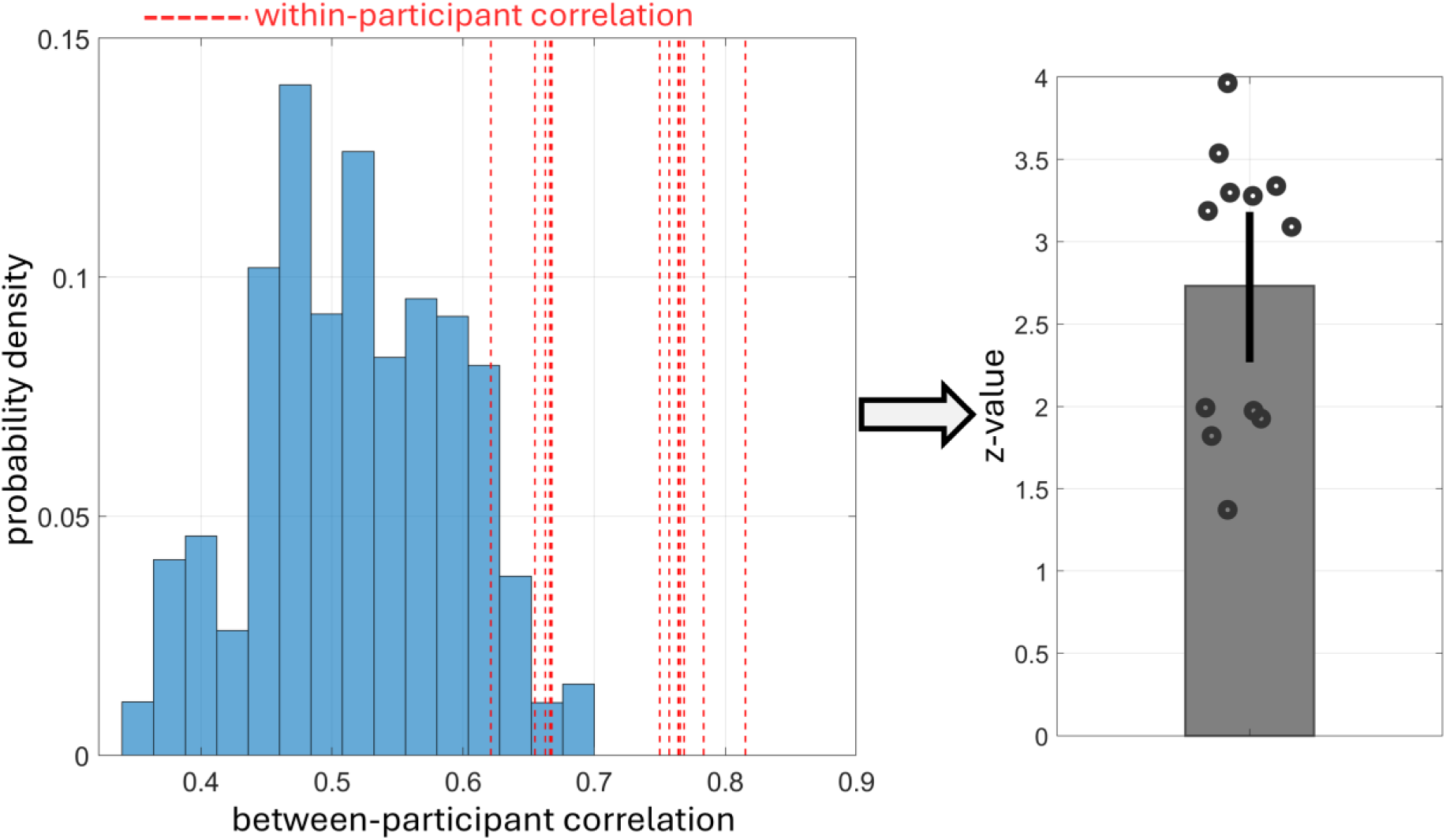
Same as Figure 2, but with the dissimilarity matrices derived using the main pre-registered approach (i.e., estimating ranking probabilities and then applying a 5D MDS to reduce dimensionality and thereby reduce noise). The dissimilarity matrices derived using the main pre-registered approach could reliably capture each participant’s specific subjective perceptual dissimilarity. This was comparable, though slightly weaker, compared to the specificity of the dissimilarity matrices derived with the alternatively pre-registered machine-learning approach (see Figure 2, where the z-value was above 2.3 in most (ten) participants).

**Supplementary Figure 8.**
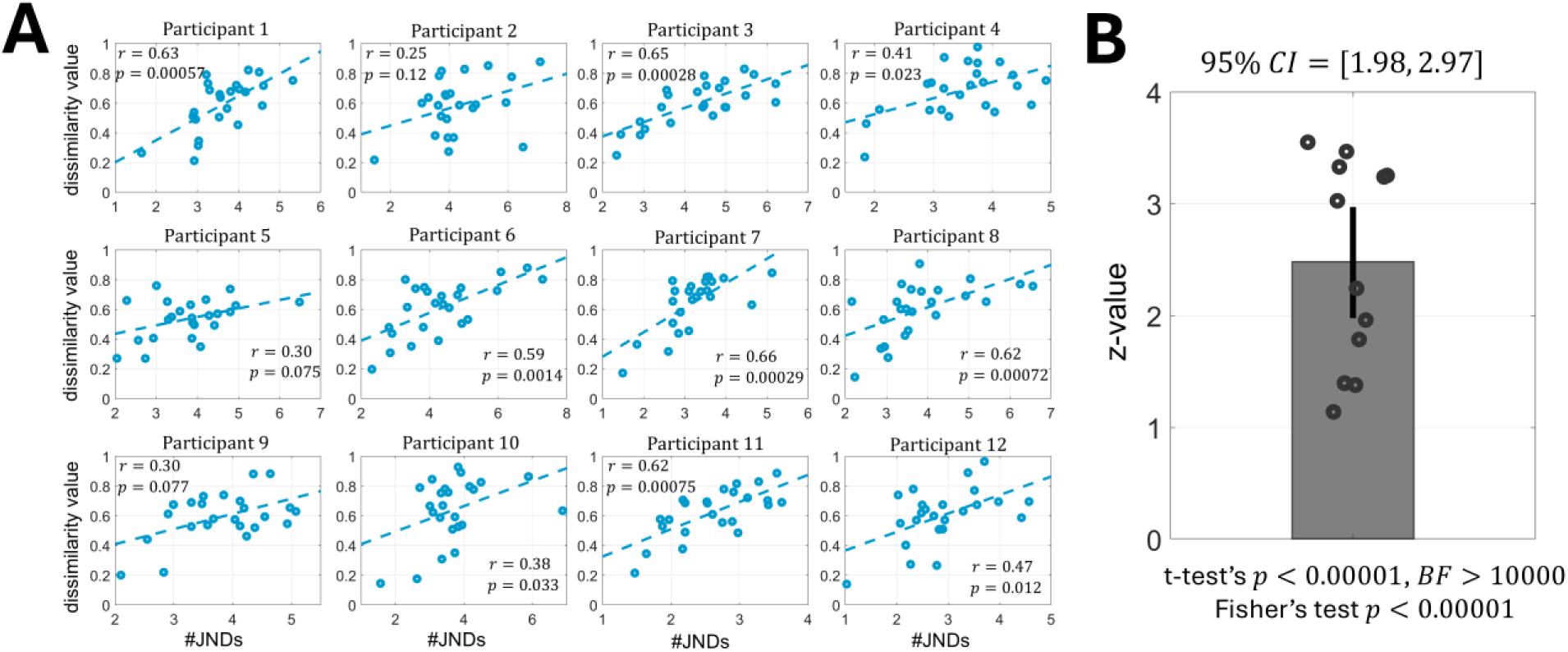
Same as Figure 3, but with the dissimilarity matrices derived using the main pre-registered approach (i.e., estimating ranking probabilities and then applying a 5D MDS to reduce dimensionality and thereby reduce noise). Similar to Figure 3, a significant positive relationship was observed between perceptual discrimination capacity and subjective perceptual dissimilarity. Note that, as shown in panel B, the 95% confidence interval was above zero, confirming Hypothesis 1, and its width was less than one, indicating that the data-collection stopping criterion was met.

**Supplementary Figure 9.**
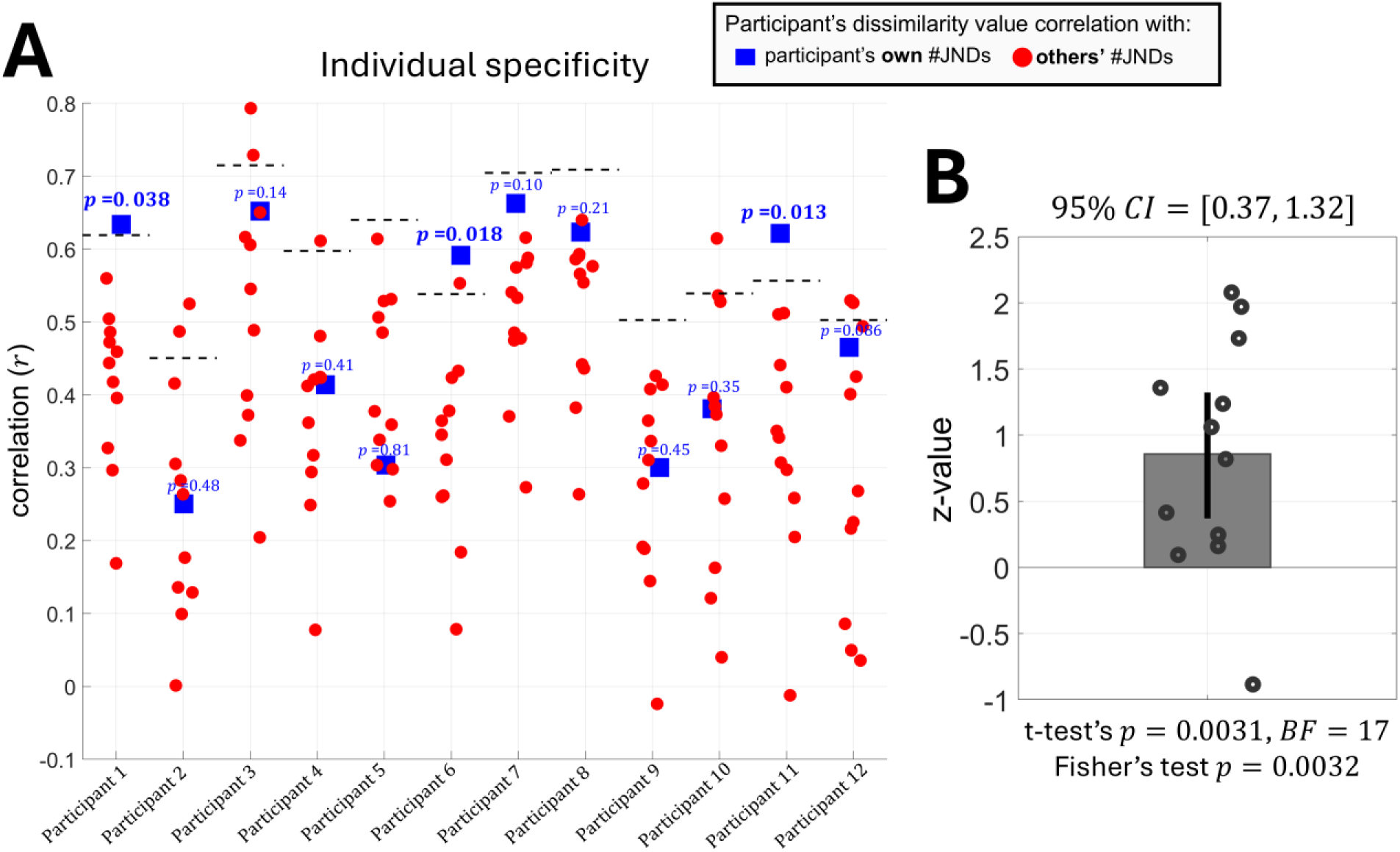
Same as Figure 4, but with the dissimilarity matrices derived using the main pre-registered approach (i.e., estimating ranking probabilities and then applying a 5D MDS to reduce dimensionality and thereby reduce noise). Similar to Figure 4, the association between perceptual discrimination capacity and subjective perceptual dissimilarity was quite specific to each individual. Note that, as shown in panel B, the 95% confidence interval was above zero, confirming Hypothesis 2, and its width was less than one, indicating that the data collection stopping criterion was met.

**Supplementary Figure 10.**
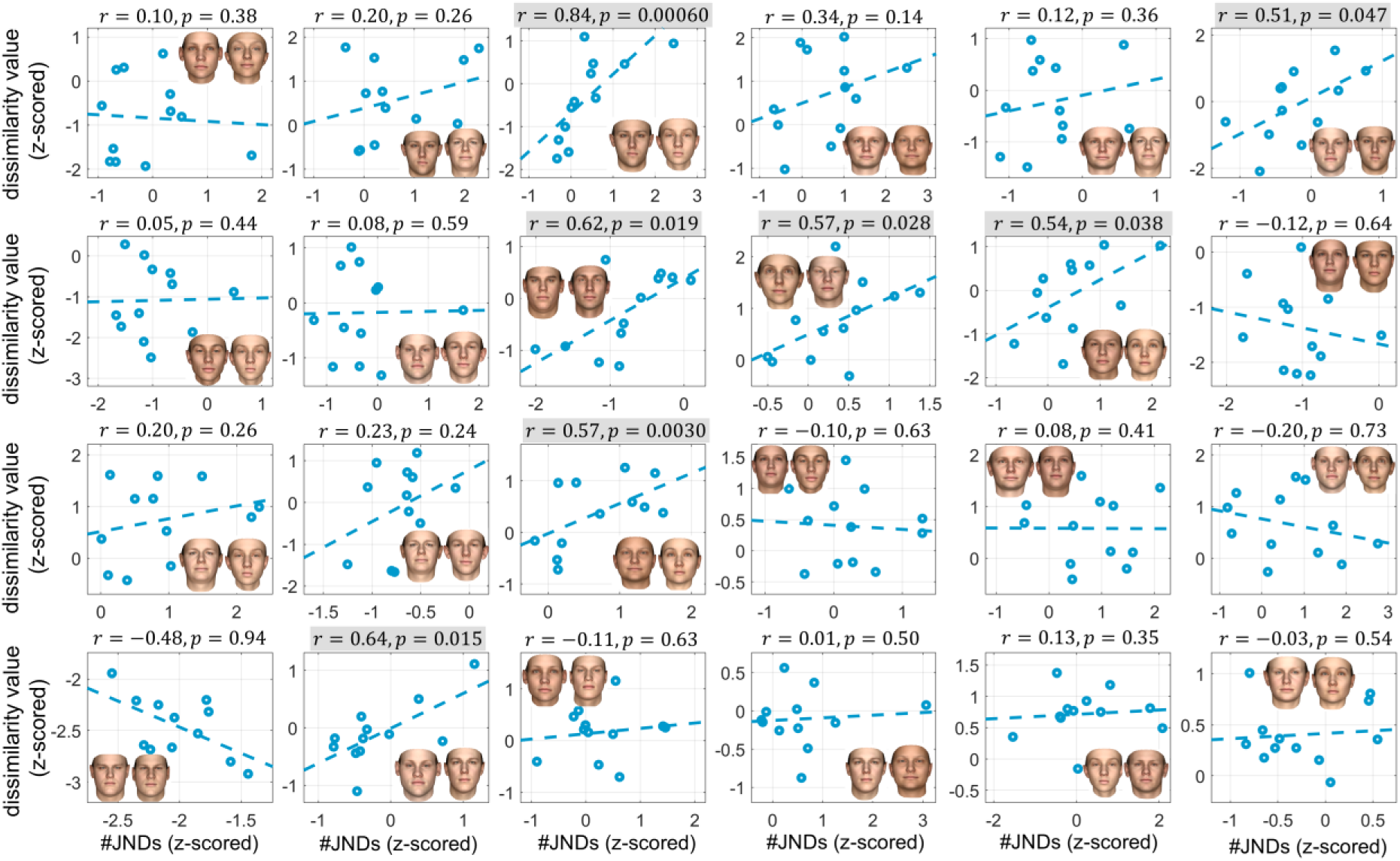
Same as Figure 5, but with the dissimilarity matrices derived using the main pre-registered approach (i.e., estimating ranking probabilities and then applying a 5D MDS to reduce dimensionality and thereby reduce noise). Similar to Figure 5, the association between perceptual discrimination capacity and subjective perceptual dissimilarity was positive in most pairs across participants, supporting our hypotheses. A t-test on the r value being positive across pairs yielded a p-value of 0.0041 and a BF of 12 (one-tailed). Also, this association was more salient in more controversial pairs (compare the last row, which shows less controversial pairs, with the other rows).

**Supplementary Figure 11.**
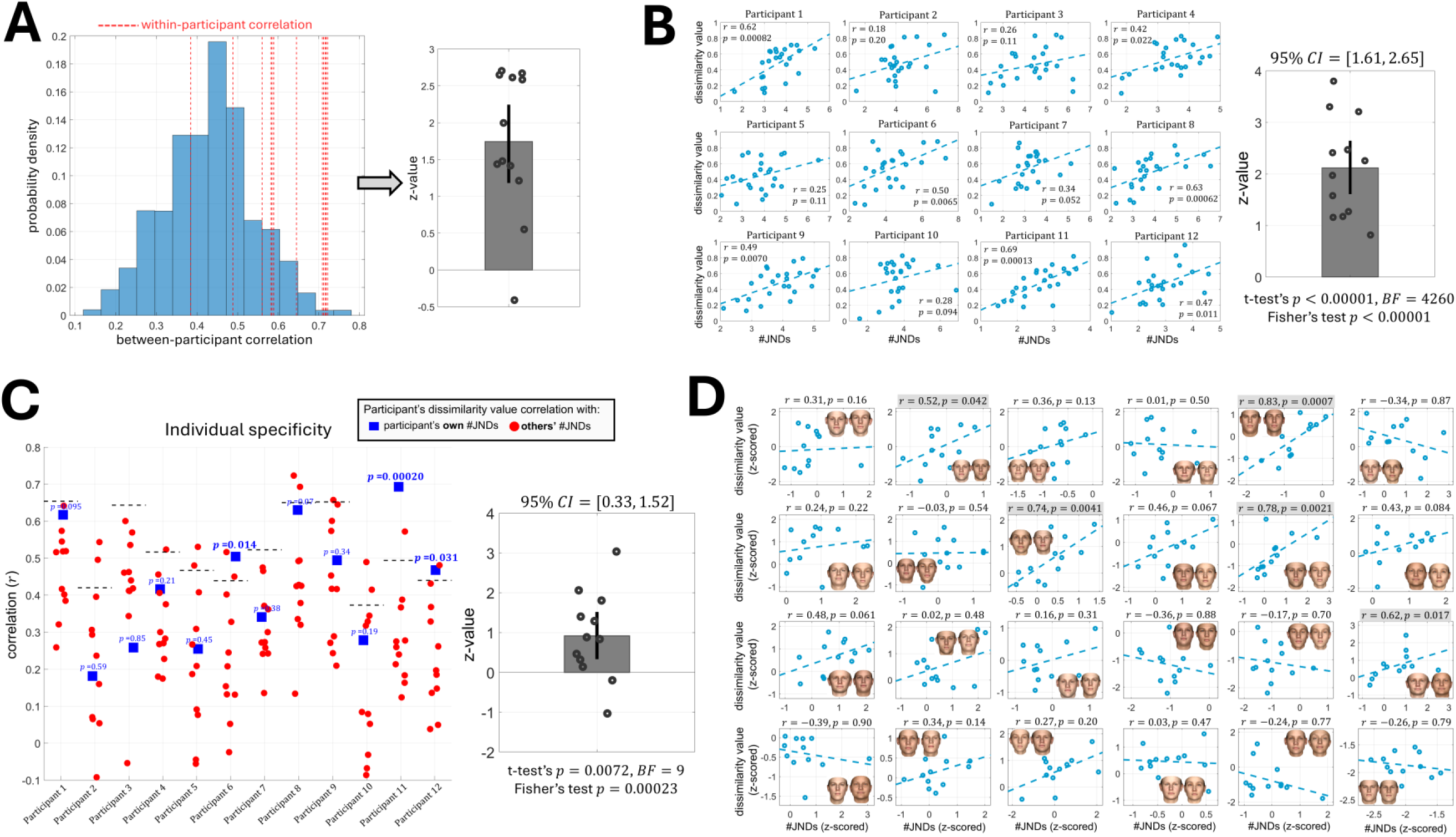
(A) same as Figure 2, (B) same as Figure 3, (C) same as Figure 4, (D) same as Figure 5, but with the dissimilarity matrices derived using the main pre-registered approach incorporating a 2D instead of a 5D MDS. Hypotheses 1 and 2 were confirmed (see panels C and D, respectively). As shown in Panel D, perceptual discrimination capacity was positively associated with subjective perceptual dissimilarity across participants in most face pairs. A t-test on the *r* value being positive across pairs yielded a p-value of 0.0077 and a BF of 7 (one-tailed). Panel A indicates that using a 2D MDS could also capture individuals’ unique subjective perceptual dissimilarity, although it performed noticeably worse than using a 5D MDS (see Supplementary Figure 7) and the alternative machine-learning approach (see Figure 2). Furthermore, the use of a 2D MDS made the relationship between perceptual discrimination capacity and subjective perceptual dissimilarity (panel B), as well as its individual-specificity (panel C), appear weaker than those observed in supplementary Figures 8 and 9 (use of a 5D MDS), and Figures 3 and 4 (use of the alternative machine-learning approach). Therefore, forcing the dissimilarity relations into two dimensions caused more harm (information loss) than benefit (e.g., noise reduction).

